# Live imaging of avian embryos revealing a new head precursor map and the role for the anterior mesendoderm in brain development

**DOI:** 10.1101/2020.08.22.262436

**Authors:** Koya Yoshihi, Kagayaki Kato, Hideaki Iida, Machiko Teramoto, Akihito Kawamura, Yusaku Watanabe, Mitsuo Nunome, Mikiharu Nakano, Yoichi Matsuda, Yuki Sato, Hidenobu Mizuno, Takuji Iwasato, Yasuo Ishii, Hisato Kondoh

**Affiliations:** Faculty of Life Sciences, Kyoto Sangyo University, Motoyama, Kamigamo, Kita-ku, Kyoto 603-8555, Japan; National Institutes of Natural Sciences, Exploratory Research Center on Life and Living Systems (ExCELLS), National Institute for Basic Biology, Okazaki, Aichi 444-8787, Japan; Institute for Protein Dynamics, Kyoto Sangyo University, Motoyama, Kamigamo, Kita-ku, Kyoto 603-8555, Japan; Avian Bioscience Research Center, Graduate School of Bioagricultural Sciences, Nagoya University, Furo-cho, Chikusa-ku, Nagoya 464-8601, Japan; Department of Anatomy and Cell Biology, Graduate School of Medical Sciences, Kyushu University, 3-1-1 Maidashi, Higashi-ku, Fukuoka 812-8582, Japan; Laboratory of Mammalian Neural Circuits, National Institute of Genetics (NIG), Mishima, Shizuoka, 411-8540, Japan; International Research Center for Medical Sciences (IRCMS), Kumamoto University; 2-2-1 Honjo, Chuo-ku, Kumamoto City 860-0811, Japan; Department of Biology, School of Medicine, Tokyo Women’s Medical University, Shinjuku-ku, Tokyo 162-8666, Japan; Institute for Comprehensive Research, Kyoto Sangyo University, Motoyama, Kamigamo, Kita-ku, Kyoto 603-8555, Japan

## Abstract

We investigated the initial stages of head development using a new method to randomly label chicken epiblast cells with enhanced green fluorescent protein, and tracking the labeled cells. This analysis was combined with grafting mCherry-expressing quail nodes, or node-derived anterior mesendoderm (AME). These live imagings provided a new conception of the cellular mechanisms regulating brain and head ectoderm development. Virtually all anterior epiblast cells are bipotent for the development into the brain or head ectoderm. Their fate depends on the positioning after converging to the AME. When two AME tissues exist following the ectopic node graft, the epiblast cells converge to the two AME positions and develop into two brain tissues. The anterior epiblast cells bear gross regionalities that already correspond to the forebrain, midbrain, and hindbrain axial levels shortly after the node is formed. Therefore, brain portions that develop with the graft-derived AME are dependent on graft positioning.

## INTRODUCTION

The primordia of embryonic neural tissues develop under distinct regulatory mechanisms between the head level (brain) and trunk level (spinal cord) (Kondoh et al., 2016). This axial level-dependent difference was first identified in our study on the regulation of the *Sox2* gene (Uchikawa et al., 2003), which indicated that *Sox2* expression in the initial step of brain precursor development is regulated by the N2 enhancer (Iwafuchi-Doi et al., 2011). In contrast, the *Sox2* expression at more posterior level involves the Wnt signaling-dependent N1 enhancer (Takemoto et al., 2006). This regional difference in the enhancer dependence of *Sox2* activation reflects the different developmental pathways of neurogenesis: The brain tissues develop directly from the epiblast in response to N2-dependent *Sox2* activation (Iwafuchi-Doi et al., 2012; Kondoh et al., 2016), whereas the spinal cord develops at least partly via the neuro-mesodermal progenitors (NMPs) (Tzouanacou et al., 2009), the dichotomic fate choice of which depends on the regulation of N1 enhancer activity, involving TBX6 in the mouse (Kondoh and Takemoto, 2012; Takemoto et al., 2011).

To investigate the details of the developmental regulation in the anterior epiblast (the epiblast roughly anterior to the node level) involved in head development, we chose chicken embryos as a model. The chicken embryos initially comprising epiblast and hypoblast layers are flat and relatively large (several mm^2^) among amniotes; they can be cultivated for a few days to support development (Chapman et al., 2001; New, 1955; Uchikawa et al., 2003). These features of early-stage chicken embryos have facilitated various experimental manipulations of the epiblast, including gene transfer by electroporation (Uchikawa et al., 2003) and grafting tissues and cells (e.g., (Dias and Schoenwolf, 1990; Linker and Stern, 2004; Storey et al., 1992)). We focused on stage 4 (st. 4) chicken embryos (Hamburger and Hamilton, 1951), a stage at which the primitive streak has grown to its maximum length with the conspicuous node at its anterior end. This is the stage when the node develops only into the anterior mesendoderm (AME) and the posterior notochord, as shown in this study.

The *Sox2* N2 enhancer activity covers the anterior epiblast at st. 4; it activates *Sox2* expression in a broad area of the anterior epiblast at st. 5 (Uchikawa et al., 2011). This observation suggests the potential of all anterior epiblast cells to express *Sox2* and hence develop into the brain. However, the presumptive brain precursor region reported in a previous study (Fernández-Garre et al., 2002) was much narrower than the region of N2 enhancer activation and confined to a node-proximal region. We considered that bridging the gap between the broad N2-activation in the epiblast and the limited distribution of brain precursors in normal embryogenesis is fundamental to understanding head development. To pursue this issue, we employed two new approaches in combination: first, to randomly label epiblast cells with fluorescence, and to live-image these fluorescent cells until the head tissues are formed; second, to examine the consequence of grafting the node or node-derived anterior mesendoderm (AME) at various positions in the anterior epiblast field by live imaging.

To randomly label chicken epiblast cells with enhanced green fluorescent protein (EGFP), we adapted the Supernova technique developed for labeling mouse neurons (Luo et al., 2016; Mizuno et al., 2014) to rapidly developing chicken embryo epiblast. The live imaging of the Supernova-driven EGFP (SN-EGFP)-labeled epiblast cells uncovered two essential steps of epiblast reorganization, leading to head development. In the first step occurring most actively between st. 5 and st. 6, the epiblast cells collectively migrate anterolaterally and converge on the head axis. This collective cell migration occurs broadly even in the distance from the periphery of the N2-positive epiblast region. However, the timing to initiate cell migration or the migration rate varied markedly among embryos and even between the sides of an embryo. These observations suggested a certain degree of stochasticity in the positioning of brain precursors in the epiblast. In the second step, the epiblast cells that converged close to the head axis start to form the neural plate. Only then will the positioning in the neural plate determine the cell’s developmental fate into brain portions, regardless of the initial positioning in the epiblast field. Likewise, cells immediately outside the neural plate develop into the overlying head ectoderm of the same axial level as adjacent brain precursors. These observations suggest that whether a cell participates in brain development or development of the overlying head ectoderm is determined only at the moment of neural plate formation

To determine to what extent the brain/head ectoderm bipotentiality holds for the epiblast, we performed the following experiments. We grafted the node or node-derived AME tissues of mCherry-transgenic Japanese quail at various positions in the anterior epiblast field of SN-EGFP-labeled chicken embryos, inside the area pellucida. This was a new challenge because earlier node graft experiments were performed by choosing graft positions at the periphery or outside the area pellucida (e.g., (Dias and Schoenwolf, 1990; Storey et al., 1992; Streit et al., 1997)). We then performed live imaging of both the graft and the epiblast. We found that the node-derived AME (that develops into the prechordal plate and anterior notochord), rather than the node, provided the second center for epiblast convergence, so that the centrally positioned epiblast cells developed into the secondary brain tissue, and surrounding cells into the associated head ectoderm.

These live imaging-based analyses provided us with a new conception of the cellular mechanisms leading to development of the brain and head ectoderm. Virtually all anterior epiblast cells have the potential to develop into the brain or overlying head ectoderm, which depends on the developmental context. When an AME develops underneath the anterior epiblast, the epiblast cells converge on the position of the AME, and the cells positioned proximal to the AME axis develop into the brain tissue. In contrast, those located more distally develop into the head ectoderm. When two AME tissues co-exist, for example, following an ectopic node graft, the epiblast cells converge on the two AME positions and develop two brain tissues. Under such conditions, the two AMEs compete for the converging epiblast cell pool. Thus, the fate of anterior epiblast cells toward the neural or head ectoderm depends on the final allocation in the AME-convergent epiblast populations. However, the anterior epiblast cells bear gross regionalities that correspond to the forebrain (FB), midbrain (MB), and hindbrain (HB) axial levels after head formation. Hence, the brain portions that develop along with the graft-derived AME (gAME) depend on the anteroposterior (AP) graft positioning. These mechanisms also account for various earlier observations and provide the foundatiosn for future investigations concerning vertebrate brain development.

## RESULTS

### Random fluorescent labeling of epiblast cells using a Supernova technique revealed the convergence of cells from a wide area toward the embryo axis

A Supernova technique was previously used to randomly label cells with fluorescent proteins in developing mouse brain for a few days (Luo et al., 2016; Mizuno et al., 2014). We adopted this technique to randomly label chicken embryo epiblast cells more quickly (Fig. 1A). In the original version of the Supernova system, tetracycline-responsive element (TRE)-*Cre* vector (b) and CAGGS-floxed Stop-EGFP-IRES-tTA vector (c) were co-introduced in mouse embryos to label neuronal clones (Luo et al., 2016; Mizuno et al., 2014) (Fig. 1A). The TRE spontaneously activates the *Cre* gene infrequently due to its leakiness, and elicits EGFP-IRES-tTA expression via removal of a floxed-Stop sequence, which feed-forwardly activates TRE-Cre to remove all floxed-Stop sequences 5′ of EGFP-IRES-tTA transgenes. To activate EGFP-IRES-tTA more quickly, we added a small amount of a tTA expression vector (Fig. 1(A)(a)) to facilitate the feed-forward EGFP-IRES-tTA activation process. This modification was successful. Following electroporation of chicken embryos of st. 4 in culture with the vector cocktail from the epiblast side (Uchikawa et al., 2003), EGFP-labeled cells became detectable usually in 1 h, increased their number, and became distributed over the epiblast area in 2 h of electroporation when the embryos were still at late st. 4 to st. 5.

**Figure 1.**
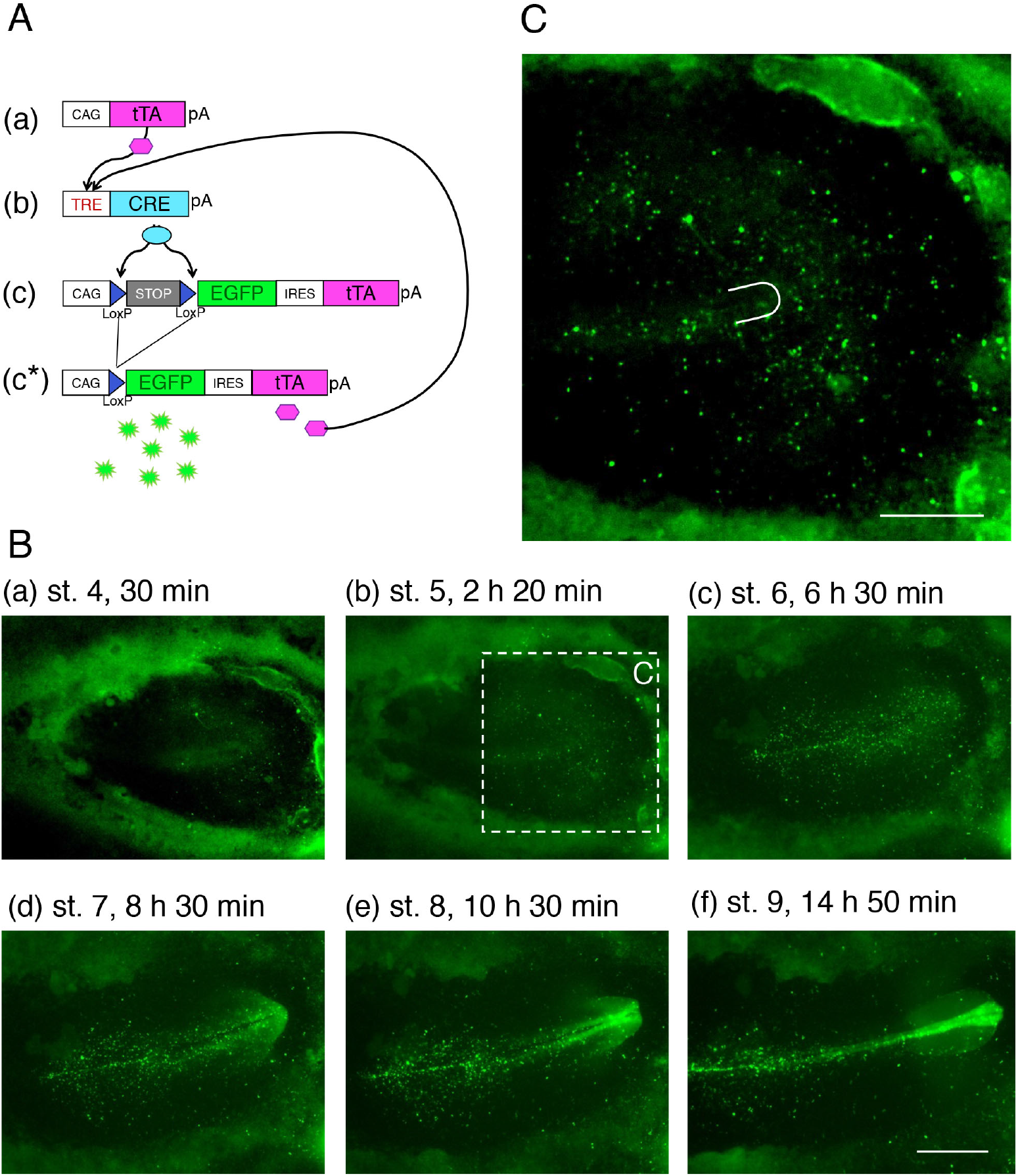
The performance of Supernova-based epiblast cell labeling with EGFP. (A) The vector composition in the Supernova system. (a) An expression vector to provide an amount of tTA [tetracycline-controlled transactivator (Gossen et al., 1995)] in electroporated cells. (b) TRE (tetracycline-responsive element)-dependent CRE expression vector. When tTA binds to this element, CRE recombinase is synthesized. (c) An EGFP-tTA joint expression system using an IRES (internal ribosome entry site) sequence driven by the CAGGS enhancer-promoter complex (CAG) (Niwa et al., 1991). This vector is delivered more abundantly than the other two in electroporated cells. However, the coding sequences remain untranscribed unless the transcriptional termination sequence cassette (STOP) flanked by the LoxP sequences is excised by CRE, supplied by the activation of the vector (b). (c*). The tTA proteins expressed from the STOP-excised vector (c) feed-forwardly activate vector (b), producing CRE recombinases more abundantly. This process continues until all vector (c) sequences become free from the STOP cassette, leading to the full EGFP expression in electroporated cells. (B) An example of Supernova-labeled chicken embryo at representative developmental stages with indications of the time that elapsed after the start of live recording. These frames are excerpts from Movie A. In (a), the EGFP signals were additionally enhanced to visualize the initially low EGFP signal. The bar indicates 1 mm. (C) An enlargement of the embryo region indicated by the broken rectangle in (B)(b) with EGFP signal enhancement as in (B)(a). A U shape indicates the node position. The bar indicates 500 μm.

A representative time-lapse recording is shown in Movie A (15 h recording with intervals of 10 min) and excerpts are shown in Figure 1(B). This embryo with electroporation of the Supernova vector cocktail mainly in the anterior part of the epiblast started to show the SN-EGFP fluorescence in 30 min of culturing (Fig. 1(B)(a)). In this and also most electroporated embryos, the density of SN-EGFP-labeled cells became sufficiently high for further analysis during the period of st. 5, as shown in Figure 1(C). Moreover, the cell displacements during st. 4 to st. 5 at levels anterior to the node were found to be small (18 ± 12 μm/h) without defined orientation) (Supplementary Figure 1). For these considerations, we analyzed Supernova-labeling data from st. 5 onward.

To characterize the migration of the SN-EGFP-labeled cells, we analyzed the trajectories of these cells. To adjust an oblique placement and time-to-time drifts of embryo images in the time-lapse recording, embryo images in all frames were reoriented so that the node position at st. 5 was kept as the posterior end of the horizontal axis of the forming head. To manage the cell-to-cell variation in fluorescence intensity and fluorescence image radius among the SN-EGFP-labeled cells, and the time-dependent changes of fluorescence intensity from the same cells, we devised a procedure to register cell positions with higher fluorescence intensity above the background. Then, we tracked the trajectories of the fluorescent points over image sequences using software developed in-house.

Figure 2(A) shows two representative cases of SN-EGFP trajectories over the period from st. 5 to 8, covering ~9 h. The embryo displayed in Figure 2(A)(a) developed a high density of labeling at epiblast regions remote from the embryo axis allowing visualization of cells migrating from a distance, whereas the embryo in Figure 2(A)(b) is the embryo shown in Figure 1, which had high-density labeling close to the embryo axis, allowing high-resolution analysis of cell migration close to the embryo axis. The entire processes of trajectory extension up to st. 9 (~15 h) are shown in Movies B1 and B2, respectively.

**Figure 2.**
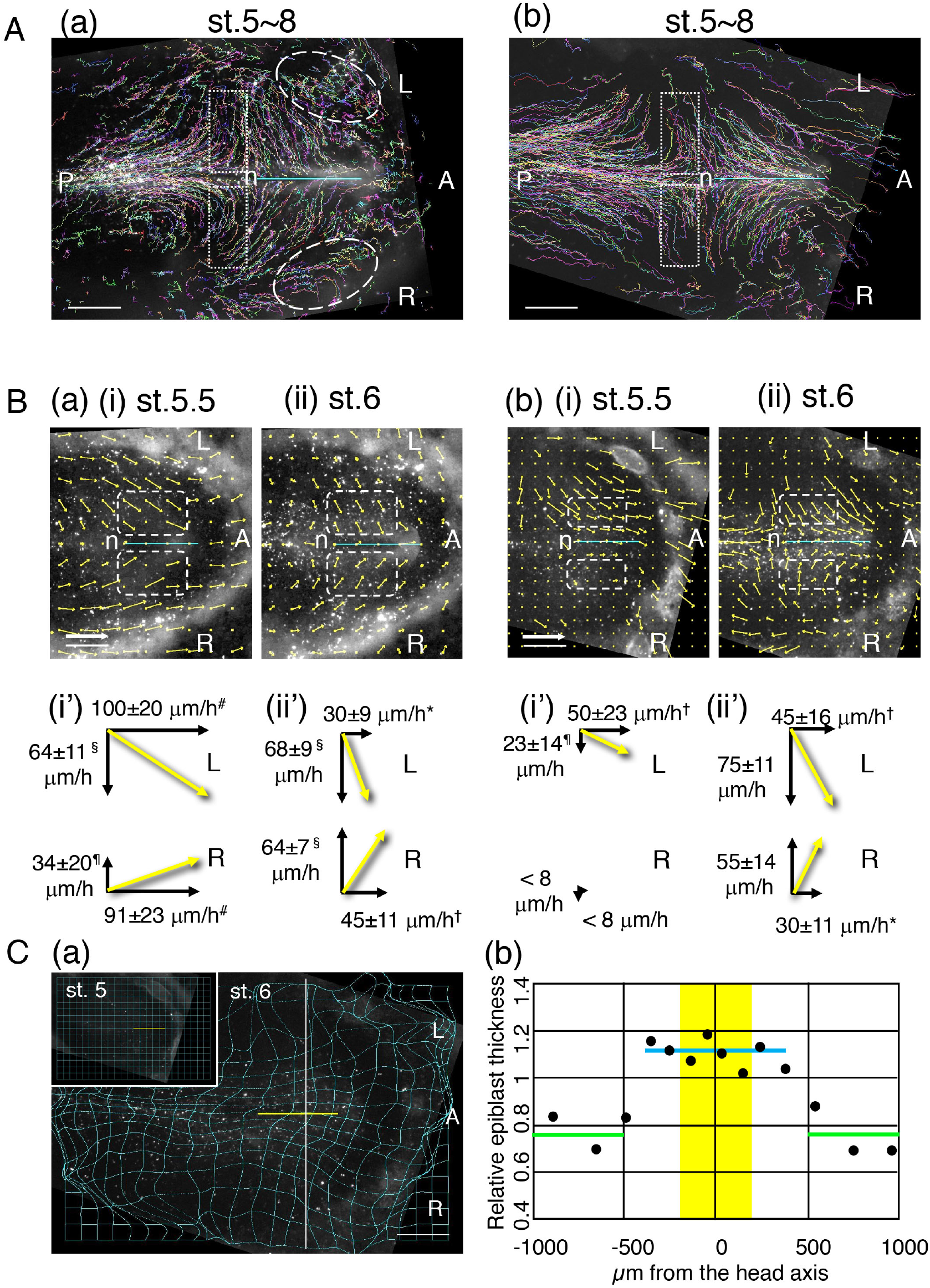
Cell tracking-based analyses of collective cell migrations that form the head tissues. (A) Trajectories of SN-EGFP-labeled cells starting from st. 5 in embryos after electroporation of SN vectors at st. 4. The cyan bars indicate the head axes, and “n” the original node position. (a) An embryo with high-density labeling in the epiblast areas distant from the brain axis. (b) An embryo with high-density labeling close to the anterior embryo axis. Snapshot data at st. 8 are shown here, whereas the full data up to st. 9 are shown in Movies B1 and B2. A, P, R, and L indicate the anterior, posterior, right, and left sides of an embryo. The broken ovals in (a) include trajectories showing cell migrations disregarding the area pellucida boundary. The broken rectangles indicate the trajectories of the cell groups that converged on the midline first and then migrated posteriorly along the embryo axis. The bars indicate 500 *μ*m. (B) Vector presentations of averaged cell migrations around the grid points on the embryo epiblast. Grid intervals were 250 *μ*m for embryo (a) and 150 *μ*m for embryo (b), reflecting labeled cell densities. The cyan bars indicate the head axes starting from the nodes (n). Snapshots of data (i) at a stage between st. 5 and st. 6 (designated as st. 5.5), and (ii) at st. 6 are presented. Overall data are shown in Movies C1 and C2. Migrations of labeled cells positioned in a grid point-centered circle with 250/150 *μ*m-radius were averaged over three consecutive frames (−10, 0, and +10 min). An incidental absence of a trackable labeled cell in the circle produced data identical to the absence of migration. Such data points [e.g., three points in the encircled areas in (a)(i)] were disregarded in the following cell migration rate calculations. Cell displacements in a 10 min frame were enlarged tenfold. An arrow size equivalent to the 500 *μ*m scale bar indicates the cell migration rate of 300 *μ*m/h. (i’) and (ii’) show averages of the cell migration vectors in the broken rectangles (covering the grid points 250 to 500 *μ*m from the head axis and 500 *μ*m anterior from the node) over three consecutive frames at st. 5.5 and st. 6, respectively. The cell migration vectors were broken down into axial (horizontal) and vertical components, and average rate in each direction with standard deviation is shown in *μ*m/h. The differences in the velocity components along or vertical to the head axis were not significant (p > 0.15) within the groups indicated by the symbols (#*†§¶); otherwise, the differences were significant (p < 0.05) (Source data 1). (C) Reorganization of the anterior epiblast sheet in the cell arrangements and regional cell densities, as indicated by the deformation of the square lattice at st. 6 starting from the square at st. 5 (inset). Data for the embryo in (B)(b) with grid intervals of 150 *μ*m were used. (a) Deformation at st. 6 of the square lattice at st. 5 (inset). The calculated grid positions at st. 6 were connected by splines to visualize the deformations. The white horizontal bar indicates 500 *μ*m. The yellow horizontal bar indicates the head axis with the posterior end at the original node position. (b) The estimated relative cell thickness (reciprocal to the lattice-surrounded areas relative to the square unit in the inset) along the vertical line in (a) plotted against the distance from the head axis, assuming invariant cell volumes. Following the medial convergence and anterior elongation of the areas, the peripheral region (≥ 500 *μ*m from the axis) became thinner (green bars), whereas ≤ 400 *μ*m from the axis, a thicker cell zone (blue bar), later developing into head tissues, was formed. The neural plate occupied the narrower central zone highlighted in yellow, 200 *μ*m bilaterally from the axis, as shown in Figure 3.

The trajectories of SN-EGFP-labeled cells indicated that the cells that were widely distributed in the epiblast at the beginning converged toward the embryo axis as a collective migration. The cells initially positioned close to the area pellucida margin migrated more than 1 mm during the 9 h. The cell migration trajectories were composites of the two components, the convergence toward the embryo axis, and movements along the embryo axis. The migration along the embryo axis was oriented anteriorly or posteriorly, depending on the axial positions relative to the node. At the axial levels immediately posterior to the node, massive cell convergence toward the embryo axis initially occurred, followed by extensive and long-distance posterior migration.

### Embryo-dependent variation in the cell migrations during developmental stages 5 to 6

To investigate the cell migrations in detail in order to resolve the direction and the rate temporally (developmental stage) and spatially (embryonic region of the stage), we processed the cell trajectory data and presented average cell movement vectors at grid points drawn on the epiblast, as shown in Figure 2(B). For embryo (a), the grids were placed at 250 μm intervals. The movement vectors of all SN-EGFP-labeled cells positioned in a circle with a radius of 250 μm centered by a grid point were averaged over the three frames (−10, 0 and +10 min) and represented by a vector arrow. An arrow length equivalent to 500 μm in the space distance indicates a movement rate of 300 μm/h or 50 μm/10 min. For embryo (b), the grid distance and circle radius were 150 μm; otherwise, the parameters were the same as for (a). The entire data are shown in Movies C1 and C2, and snapshot data for a stage between st. 5 and 6 (designated as st. 5.5) and at st. 6 are presented.

The overall analysis of movement vectors indicated that the most extensive cell migrations in anterior and axial directions occurred between stages 5 and 6, a period spanning 4 to 5 h. After stage 6, when the neural plate was formed, the anterior movement of cells outside the neural plate was reduced (Movies C1 and C2).

The average migration rates in the lateral areas along the head axis indicated by broken rectangles in Figure 2(B) were estimated at respective developmental stages. The average cell migration rates were decomposed into the axial and perpendicular components are displayed in the lower panels in Figure 2(B).

These analyses indicated that the patterns of collective cell migrations were highly variable among embryos and even between the sides of embryos. Around the middle of st. 5 and st. 6 [st. 5.5 in Fig. 2(B)], cell migration rates in either the anterior or the axial direction in an embryo (a) were almost twice those on the left side of the embryo (b). The initiation of cell migration on the embryo (b) right side was delayed until st. 5.5 had passed. This delay was confirmed by inspecting the original trajectory Movie B2. However, the cell migration rates after st. 6 became similar among embryos and between embryo sides, and embryos developed normally regardless of the cell migration profiles between st. 5 and st. 6. These developmental stage-dependent changes in the collective cell migration were confirmed in similar SN-EGFP-labeled embryos (N = 8; 4 trajectory analyses, 4 visual inspections of movie data).

The embryo axis-directed collective cell convergence on a plane predicts the relative thickening of the cell layer close to the axis relative to that in the surroundings, assuming that the cellular volumes are maintained during the process. The reorganization of the SN-EGFP-labeled anterior epiblast sheet, during the period from st. 5 to st. 6, was investigated by examining the deformation of the initially square grids on the sheet (Fig. 2(C)(a)). This square grid deformation analysis was performed using the embryo shown in Figure 2(B)(b). The head axis-converging epiblast cell movements, concomitant with anterior cell migration, as indicated in Figure 2(A)(b), were reflected by the anteriorly elongating square deformation (Fig. 2(C)(a)). This convergence resulted in the thinning of the peripheral region ≥500 μm from the axis (Fig. 2(C)(b)). In contrast, a thicker epiblast zone was formed ≤400 μm from the axis. As shown in Figure 3, only the cells within 200 μm from the axis later developed into brain tissues. Therefore, the thick medial zone presumably consists of the brain and head ectoderm precursors (Fig. 2(C)(b)).

**Figure 3.**
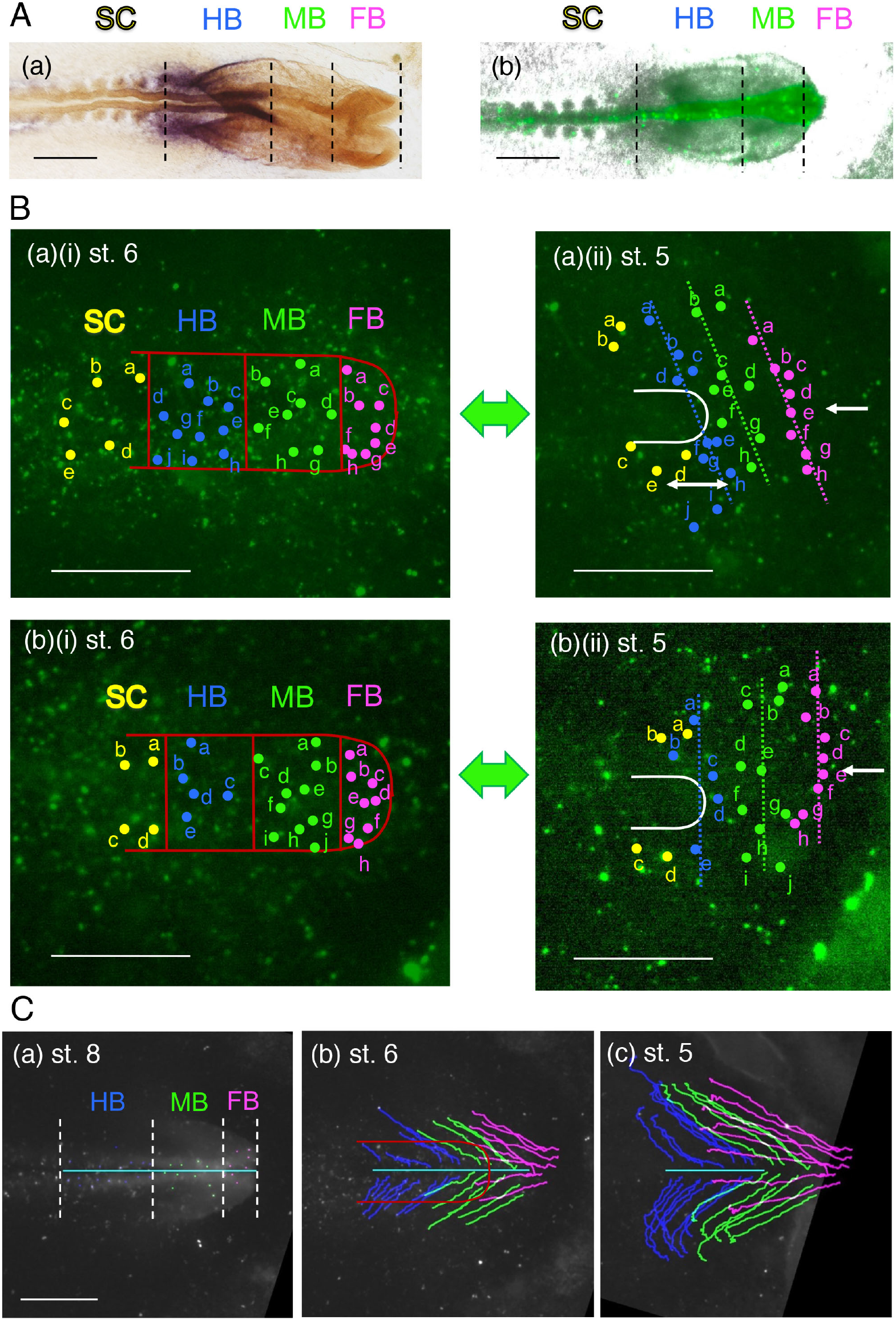
Tracking the origin of head tissue portions at different axial levels back to st. 5. (A) Determination of the brain portions in the SN-EGFP-labeled embryo at the endpoint of live imaging. (a) An embryo fixed at st. 9 when the live imaging was terminated and hybridized for *Gbx2* (brownish purple) to determine the hindbrain (HB) portion and for *Otx2* (orange) to determine the midbrain (MB) and forebrain (FB) portions. The brain portion boundaries for HB/MB and HB/spinal cord (SC) were determined by the anterior and posterior ends of the *Gbx2*-hybridized region. The FB/MB boundary was determined as the posterior end of the FB lateral bulging in the flat-mounted specimen. (b) The endpoint SN-EGFP image of the embryo superimposed on the bright-field image. This image was aligned with the in situ hybridization image in (a). The brain portion boundaries were traced back to st. 8 (C) and st. 6 (B), using brightly labeled landmark cells as guides. (B) The distribution of the precursors for individual brain portions of two embryos at st. 6 (a)(i) and (b)(i) and st. 5 (a)(ii) and (b)(ii). Embryo (a) was the same embryo as shown in (A). The precursor regions for individual brain portions, FB, MB, and HB at st. 6. Dark red lines indicate their boundaries and the HB/anterior SC boundary. An approximate lateral limit of the brain precursors at st. 6 was 200 μm from the head axis, and the anterior limit was immediately posterior to the forming anterior ectodermal ridge. Panels (a)(ii) and (b)(ii) indicate the original positions at st. 5 of the cells in st. 6 in (a)(i) and (b)(i). The correspondences between st. 6 and st. 5 are indicated by alphabetical labeling and color-coding individual cells: magenta for FB, green for MB, blue for HB, and yellow for anterior SC. Note that the distribution of original precursor positions at st. 5 was asymmetric, in the AP levels (a) or lateral distributions (b). (C) Long tracks of SN-EGFP-labeled cells located on the brain image (representing dorsal brain cells or overlying head ectoderm) or the brain-proximal head ectoderm at st. 8 to st. 5, with color-coding of the axial levels of brain portions. The data for the embryo shown in (B)(a) are presented. Full tracking data are shown in Movie D1 and those for the second embryo in D2. (a) The selected SN-EGFP-labeled cells at st. 8. (b) An intermediate of backtracking at st. 6 is presented. The track lines with their ends within the neural plate region (marked by the dark red line) represent dorsal brain cells. (c) Track lines from st. 8 to st. 5. Are presented. Scale bars in all panels indicate 500 μm.

### Distribution of precursors for the head tissues of different axial levels at st. 4/5

To determine from where in the anterior epiblast, the individual brain portions and abutting head ectoderm are derived, we performed the following analyses. Immediately after the time-lapse recording (usually around st. 9), SN-EGFP-labeled embryos were fixed and hybridized in situ for *Otx2* (expressed in the FB and MB, stained orange) and *Gbx2* (expressed in the HB, stained brownish purple). The hybridization patterns determined the embryo brain portions at the developmental endpoint [Fig. 3(A)(a)]. The FB/MB boundary was determined as the level where the FB bulging begins. Then these boundaries were copied on the SN-EGFP image of the same embryo [Fig. 3(A)(b)] and traced back using brightly labeled cells as landmarks to st. 8 when epiblast convergence toward the embryo axis was diminished, to st. 6 when the extensive migration of the anterior epiblast was off the peak, and to the initial st. 5.

In the forming neural plate of st. 6 embryos, the position immediately posterior to the forming anterior ectodermal ridge was determined as the FB portion anterior limit, in addition to the FB/MB and MB/HB boundaries determined by the SN-EGFP back tracing. The lateral limit of the brain tissue primordium was determined as 200 μm from the embryo axis after testing forward tracking of the labeled cells. All SN-EGFP-labeled cells in the brain primordium domain, thus determined, were later incorporated in the brain tissues. A previous study using isolated grafts of neural plate portions indicated that AP patterning of the neural plate has been established by st. 6 to 7 (Rowan et al., 1999). The SN-EGFP-labeled cells into the st. 6 neural plates in two representative embryos are shown in Figure 3 (B)(a) and (b) with color-coded brain portions.

The SN-EGFP-labeled cells serving as the precursors for different brain portions were individually traced back to st. 5, to investigate the distribution of precursors at this stage. Those SN-EGFP-labeled cells successfully traced to st. 5 are shown in (a)(ii) and (b)(ii), using alphabetically labeled color codes. We found that the positioning of a brain precursor relative to the adjacent SN-EGFP-labeled precursor at st. 5, concerning AP and lateral relationships, was more- or- less conserved at st. 6, confirming the occurrence of collective cell migration during the stage span. However, the brain precursor distributions at st. 5 varied between the embryos and even between the situs. The lateral distribution of the brain precursors at st. 5 was 770 μm in the embryo (a), whereas that in (b) was narrower, 644 μm. The center of mass of the lateral distribution in embryo (a) [Fig. 3B(a)(ii) arrow] was 20 μm to the left, fairly close to the midline, whereas that in the embryo (b) was 112 μm to the left [Fig. 3B(b)(ii) arrow], indicating strong asymmetry in the distribution. Significantly, the head precursor distribution of the embryo (a) at st. 5 was arranged along the oblique lines. This observation indicated that a brain precursor on the right side 250 μm away from the axis contributed to the same brain domain as a precursor of the left side at ~200 μm posterior (the double arrow).

These variations in the brain precursor distribution are accounted for by the variations in the convergence of epiblast cells shown in Figure 2. For instance, the embryo (a) in Figure 3(B) is the same as the one in Figure 2(A)(b), which showed minimum anterior cell migration on the right side between st. 5 and st. 6 [Fig. 2(B)(b)]. As a rough estimate, a 50 μm/h difference in the anterior cell migration rates between the situs over 4 h accounts for the ~200 μm axial level difference observed in Figure 3(B)(a)(ii))(double arrow). Consequently, the left side epiblast cells at the same axial level as the FB precursors on the right at st. 5 (e.g., those labeled “g” and “h” in magenta) developed into non-neural ectoderm. In other words, the contribution of an epiblast cell to the brain or head ectoderm, or the contribution to which brain portion is determined at the cell positioning only at st. 6.

A highly asymmetric brain precursor distribution at st. 5 observed in the embryo (b) is accounted for by the overwhelming cell migration from the left side before st. 6. This embryo case confirmed that the epiblast cell fates concerning the brain or head ectoderm are not yet determined at st. 5. The data obtained from four embryos, including (a) and (b), were compiled into an overlay, as shown in Figure 4(A).

**Figure 4.**
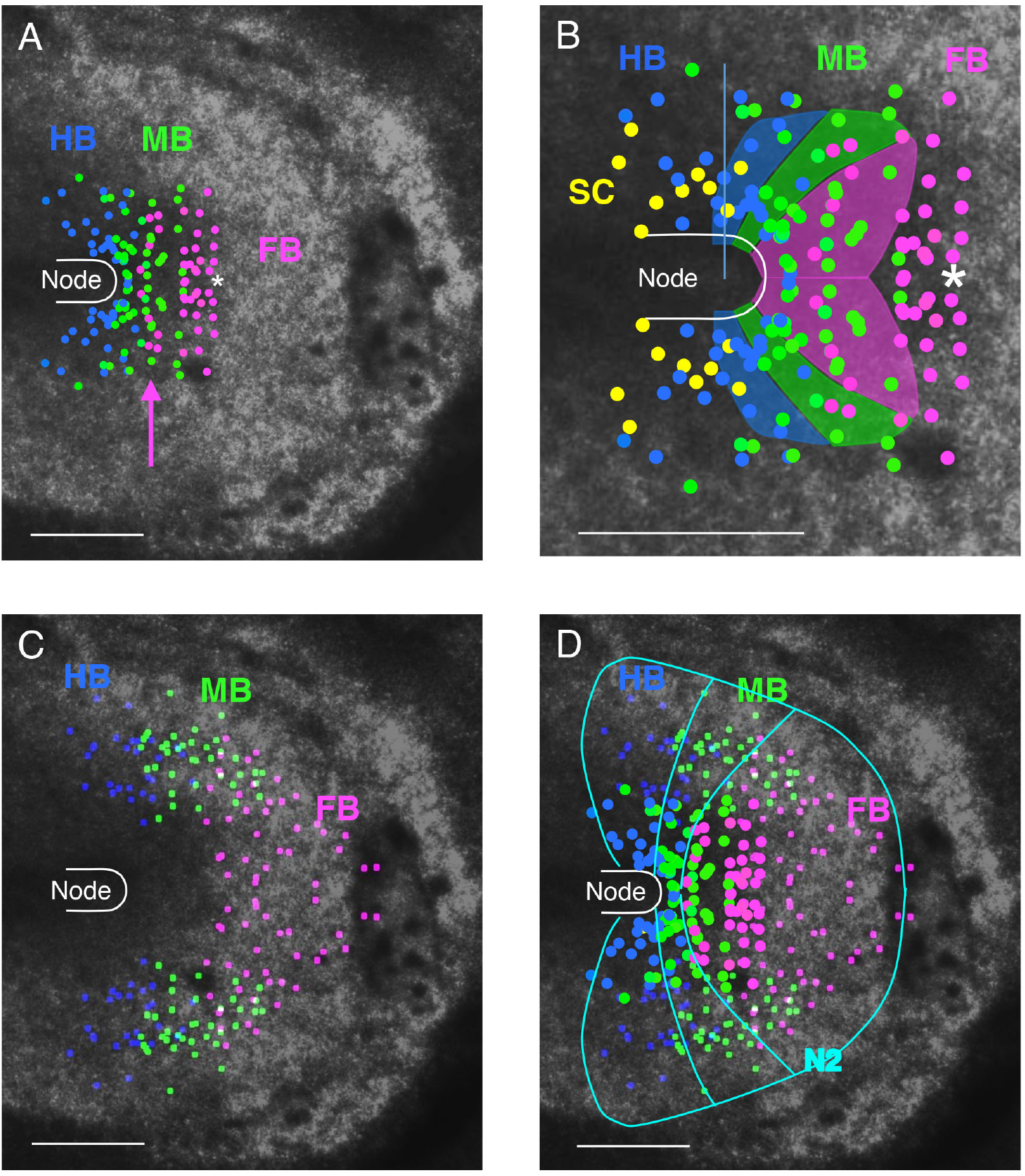
Comparison of the distribution of head tissue precursors at st. 4/5. Data for st. 5 embryos are shown. However, as the cell displacements in the anterior epiblast are minimal between st. 4 and st. 5 (Supplementary Figure 1(A)), their distributions at st. 4 are very similar. (A) Positions at st. 5 of the precursors for different brain portions. Data for embryos in Fig. 3(B)(a)(b), and two additional embryos [Supplementary Fig. 1(B)] were superimposed, which was followed by the reverse imposition. FB, forebrain; MB, midbrain; HB, hindbrain. (B) Comparison of the enlarged single-cell analysis data in (A) with the brain precursor region map drawn by Fernández-Garre et al. (2002), shown using the same color code for the brain portions. The distribution of anterior spinal cord precursors (SC) is also included to aid comparison with the data of Fernández-Garre et al. (2002). The asterisk indicates the embryo position reported to give rise to both the FB and the proximal head ectoderm (Schoenwolf and Alvarez, 1991). (C) The distribution at st. 5 of the precursors for dorsal brain/proximal head ectoderm at st. 8/9, drawn using the superimposition plus reverse imposition of three different embryo data including Fig. 3(C). (D) Combination of the data in (A) and (C) where the dots for brain precursor positions in (A) were enlarged to distinguish them from the precursors for the dorsal brain/proximal head ectoderm. The precursor regions for brain portions continue seamlessly into those of the proximal head ectoderm of the same AP levels. Rough outlines of the precursor domains are drawn in cyan lines, together with the outer limit of *Sox2* N2 enhancer activity at st. 5, taken from data of Movie E. The bars indicate 500 μm.

The undetermined brain/head ectoderm fates at st. 5 indicated that the cells developing into brain portions and those developing into proximal head ectoderm are derived from a pool of similarly positioned cells at st. 5. We performed a long-distance tracking of SN-EGFP-labeled cells of the dorsal brain and proximal head ectoderm in embryos of st. 8 to 9 back to st. 5 (a period of 14–16 h, using a modified protocol as described in Materials and Methods). Figure 3(C) shows the case of the embryo (a) starting from st. 8. Until back to st. 6, the tracking lines were arranged grossly symmetrical but became asymmetrical toward st. 5. The lines with the tracking termini in the brain portion at st. 6 (Fig. 3(C)(b)) represent cells that developed into the dorsal brain, and others to the head ectoderm. By extension of the backtracking to st. 5, many trajectories ended > 500 μm distant from the head axis. The cumulative precursor map for 3 embryos with successful long-distance tracking was drawn for dorsal brain and adjacent head ectodermal cells with the distinction of axial levels of FB, MB, and HB, as shown in Figure 4(C).

It should be noted that the st. 5 precursor positions exemplified by Figure 3(C) and cumulated in Figure 4(C) concern only the dorsal brain and the proximal head ectoderm. Comparing them with data in Figure 2(A) indicated that precursors for more lateral head ectoderm, possibly sharing the same AP regionalities, are distributed more distantly from the head axis.

We compared the distribution of the precursors for brain portions and the head ectoderm in Figure 4. The data for the distribution of the precursors for brain portions for two embryos shown in Figure 3(B) and two additional embryos [Supplementary Figure 1(B)] were accumulated, reverse-imposed, and shown in Figure 4(A). The embryo shown in Supplementary Figure 1(B)(b) provided the case with node-proximal positioning of the FB precursors (magenta arrow). The FB, MB, and HB precursors were arranged along the AP axis in the epiblast region from ~500 μm anterior to the node and covering a width of ~950 μm. The overlap of the region for precursors for different brain portions reflects the variability in the distribution among embryos. We collected data for st. 5 embryos, but the brain portion precursor distribution is considered to be similar at st. 4, given the minimal cell displacements between st. 4 and st. 5 in the anterior epiblast [Supplementary Figure 1(A)].

These data were compared with the previously published brain precursor map at st. 4 (Fernández-Garre et al., 2002) using enlarged panels in Figure 4 (B). This map of Fernández-Garre et al. (2002) was based on the homotypic transplantation of labeled epiblast plugs of ~125 μm diameter. The boundary lines were drawn on the assumption that all boundaries between precursors for brain portions and between brain precursors and head ectoderm are already determined by st. 4, an assumption not supported by our data shown in Figure 3(B). Considering these points, our data and those of Fernández-Garre et al. (2002) are generally consistent. The only significant difference was the anterior limit of ~500 μm from the node in our data for the FB precursor positioning in contrast to ~250 μm, according to Fernández-Garre et al. (2002). It should be noted that Schoenwolf and Alvarez (1991) reported that the plug grafts at the position indicated by an asterisk in Figure 4(B) produced both the FB and adjacent head ectoderm, consistent with our data. Possibly, the embryos used by Fernández-Garre et al. (2002) were those showing more extensive anterior migration between st. 5 and st. 6, resulting in the FB precursor positioning closer to the node than in some of our embryos and those used by Schoenwolf and Alvarez (1991).

The distribution of the precursors for the dorsal brain and proximal head ectoderm determined by the procedure shown in Figure 3(C) is drawn in Figure 4(C) as cumulative data of three embryos. The distribution covered the wide epiblast area in the way surrounding the brain portion precursor region [Fig. 4(A)], yet with significant overlap at their margins. Schoenwolf and Alvarez (1991) already reported the contribution of the peripheral epiblast cells to the brain-abutting head ectoderm. Precursors for a brain portion and those for dorsal brain/head ectoderm of the same AP level were arranged in continuing territories. Superimposition of these precursor distributions (distinguished by the dot size) in Figure 4(D) confirmed the seamless extension of the brain portion precursor regions to those of head ectoderm precursors at the same AP levels. All of these precursors were within the N2 enhancer activation domain indicated by the outer cyan line.

### Characterization of the st. 4 node

The analysis delineated in Figure 3(B) suggested that the brain precursors at st. 5 are bipotential, at least concerning later development into the brain portion, or the adjacent head ectoderm. The immediate question is to what extent the bipotential nature of st. 4/5 precursors extends to the outer zone precursors, which develop into the head ectoderm in normal embryogenesis. To address this issue, we planned a series of experiments to graft the second node in the outer zone. After initial trials, we realized that the strictly st. 4 of the node donor embryo is critical for eliciting the secondary head tissue development. For this reason, we characterized the st. 4 nodes first.

The stage 4 embryos, the stage usually reached after 16–20 h of incubation of eggs, are distinguished from those of other stages by the presence of a central depression of the primitive streak, thickening of the anterior end of the node, and the absence of anterior mesendoderm (anterior notochord and prechordal plate) (Hamburger and Hamilton, 1951). It is only the node of this stage at which grafting in a host embryo results in secondary brain tissue development (Dias and Schoenwolf, 1990; Storey et al., 1992). To characterize the cellular events that occur following node grafting, we excised the nodes of st. 3 to st. 5 Japanese quail embryos, grafted them in stage-matched host chicken embryos replacing the host node (N ≥ 2 at each stage) [Fig. 5(A)] [procedures are shown in Supplementary Figure 2(A)], and cultured the node-grafted embryos. After 18 h of culture, the embryos were fixed for whole-mount staining with anti-quail antibody (QCPN) to visualize node graft-derived tissues in host embryos (magenta), and for SOX2 expression to mark neural tissues (green) [Fig. 5(B)].

**Figure 5.**
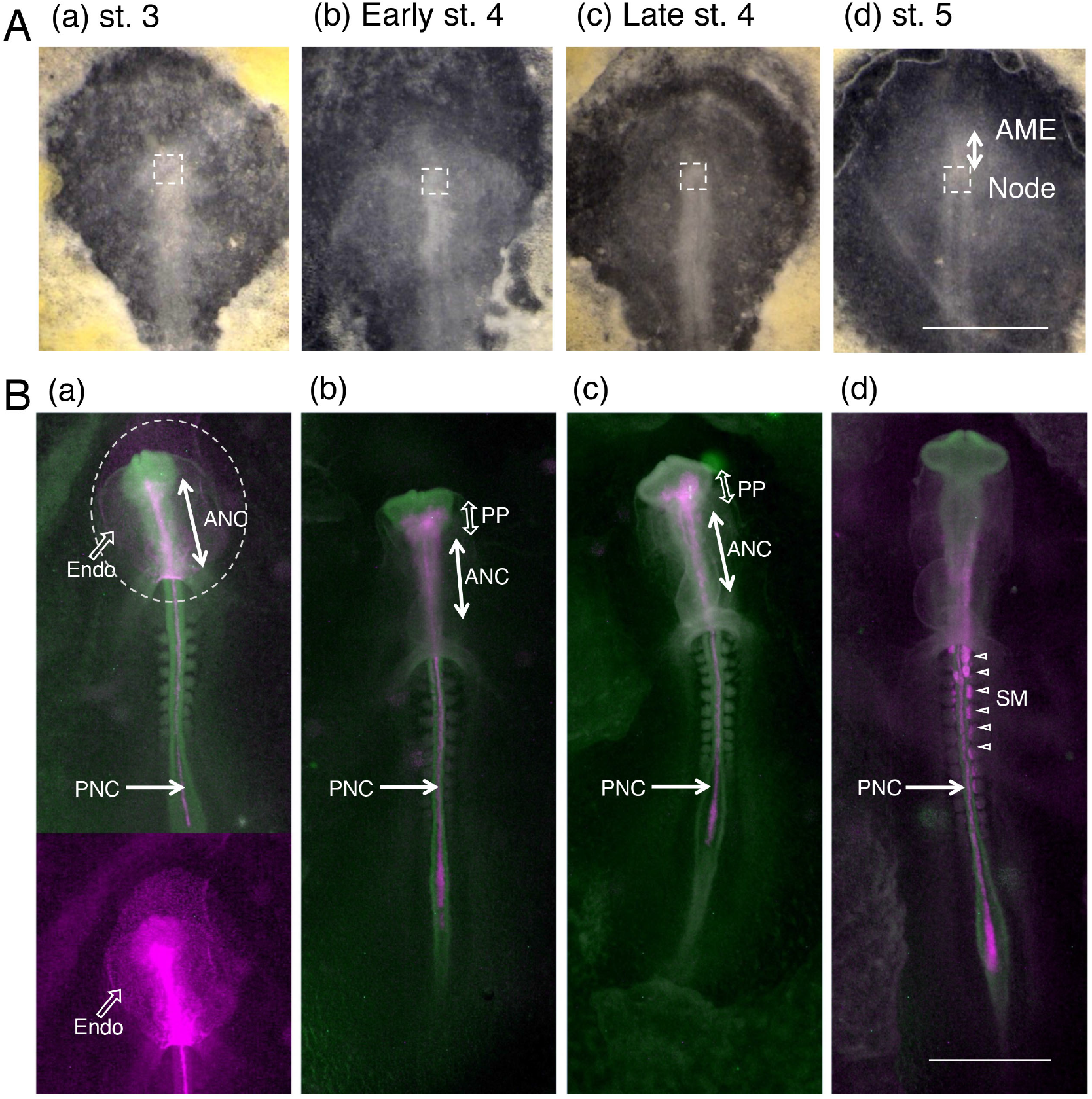
Development of st. 4 nodes solely into the anterior mesendoderm and posterior notochord. (A) The Japanese quail donor embryos in the ventral view at respective stages, and the node tissues to be excised indicated by the broken rectangles. In the st. 5 embryo the anterior mesendoderm (AME) is visible as the anterior tissue protrusion from the node and is indicated by the double arrow. The bar indicates 1 mm. The anterior is at the top. (B) The host chicken embryos that received the respective node grafts. The embryos were cultured for ~16 h, fixed, immuno-stained in whole-mount for quail-derived tissues (magenta) and SOX2 expression (green). Ventral views are shown. (a) An embryo that received a st. 3 node graft. The grafted node developed into the endoderm (Endo), encircled by the broken line, the anterior notochord (ANC), and the posterior notochord (PNC, indicated by an arrow). The lower panel shows only the anti-quail immunofluorescence to more clearly indicate development of the quail node-derived endoderm. (N = 2). (b) and (c). Embryos that received Japanese quail node grafts of early st. 4 (N = 3) and late st. 4 (N = 2). In both cases, the grafted node developed into full AME consisting of the prechordal plate (PP) and ANC, besides PNC. (d) An embryo that received a st. 5 node. (N = 3). The grafted node did not give rise to any AME tissue, but developed into the medial portions of the somites (SM with open arrowhead), besides PNC. The bar indicates 1 mm.

The st. 3 nodes developed into endoderm (Endo) [Fig. 5(B)(a)], the encircled region), the AME consisting primarily of the anterior notochord (ANC), and the posterior notochord (PNC). The nodes at st. 4 [Fig. 5(B)(b)(c)] developed only into the full AME consisting of the prechordal plate (PP) and ANC, in addition to PNC during this developmental stage. In contrast, st. 5 nodes did not produce AME, but self-differentiated into the medial portion of the somites (SM), besides PNC [Fig. 5(B)(d)]. To test the developmental potential of the anterior epiblast, we only used st. 4 node graft, which produced neither graft-derived endoderm nor somites.

The development of the st. 4 nodes was investigated by live imaging using the node of mCherry-transgenic Japanese quail (Huss et al., 2015) grafted in an SN-EGFP-labeled chicken embryo (N = 2). The grafted node fused to the host tissues in 1 h, first extended anteriorly to produce the AME that further developed into the PP and ANC, and second extended the PP (Supplementary Figure 3, Movie F). This observation indicated that although ANC and PNC appear to be a continuous single tissue, their development is temporally distinct.

### AP level-dependent development of the st. 4 quail node graft and host-derived secondary brain portions

We investigated the consequences of grafting st. 4 Japanese quail nodes at various positions of st. 4 chicken embryos mostly outside the brain precursor region (Fig. 6A). After culturing for 16–18 h, when the embryo developed to st. 9 to 10, the embryos were fixed and stained for quail-derived tissues and SOX2. Figure 6(B) displays embryos with the node graft positions indicated in Figure 6(A). The outcomes differed depending on whether the node grafts were performed on anterior or posterior epiblast.

**Figure 6.**
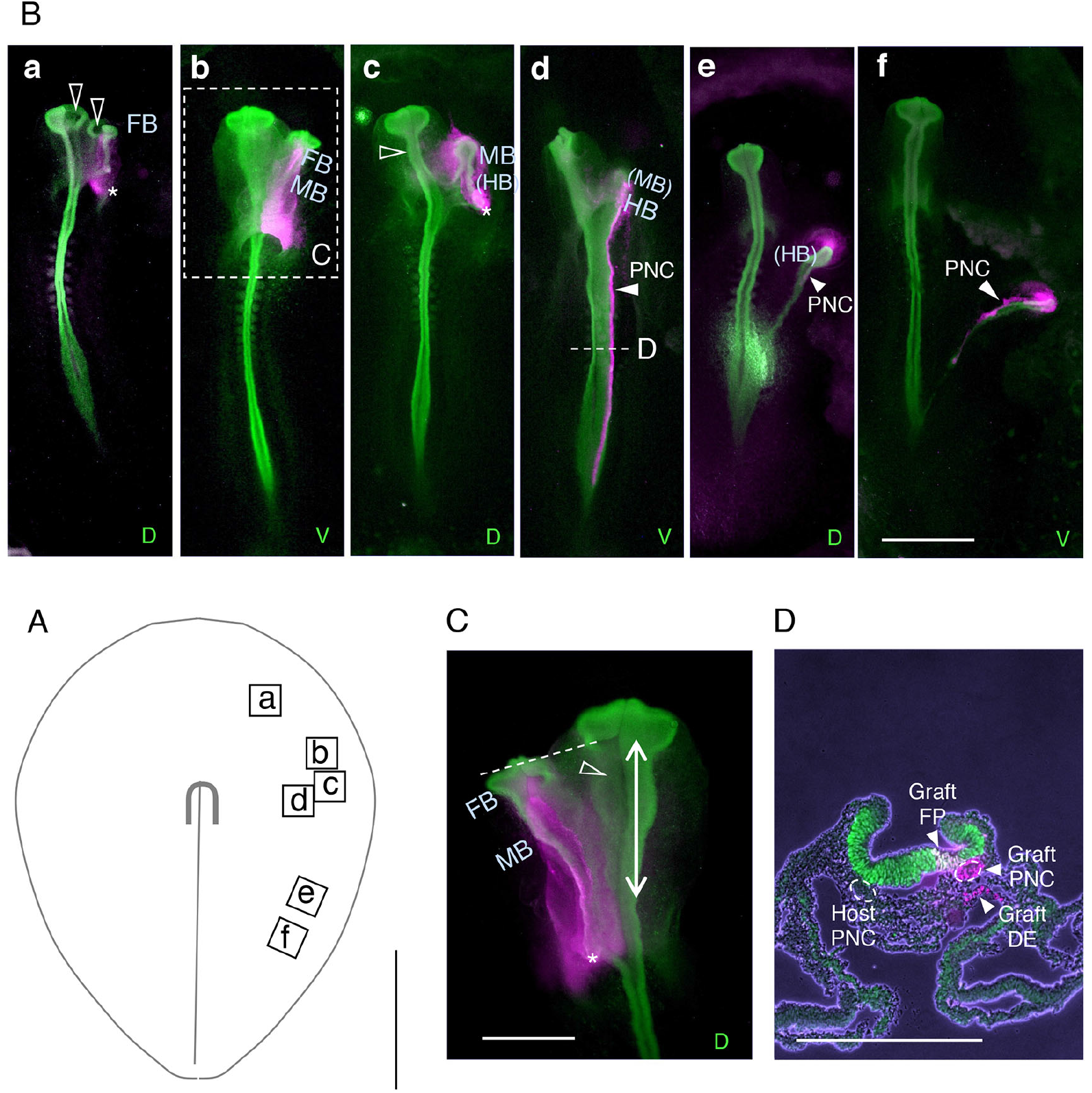
AP level-dependent development of the st. 4 quail node graft and host-derived secondary brain portions. (A) The positions of grafting st. 4 Japanese quail nodes (squares) relative to the chicken host node. The anterior is at the top. The bar indicates 1 mm, Because the zona pellucida boundaries vary among embryos, these graft positions do not indicate any spatial relationship with the zona pellucida contour schematically drawn by the thin line. The quail nodes were grafted on either side but drawn on the right side in the scheme to indicate the anteroposterior levels. (B) The node-grafted embryos after incubation for 16-20 h, immuno-stained for SOX2 (green) and quail-derived tissues (magenta). D, dorsal images; V, ventral images. The secondary brain portions judged by their morphology are labeled as FB, MB, and HB. A parenthesized brain portion indicates a minor contribution. The bar indicates 1 mm. (a) The node graft at the most anterior level resulted in the secondary FB development. Note that the dorsal facing sides of the primary and secondary forebrain portions were missing (open arrowheads). (b) The secondary brain consisted of posterior FB and MB. The enlarged dorsal view of the area indicated by the broken rectangle is shown in panel (C). (c) An embryo with a secondary brain consisting of isolated MB and a small HB. The midbrain level of the primary brain became very thin (open arrowhead). The posterior side of the graft-derived tissue ended as a cell clump, possibly a remnant of PNC. (d) The case where the node graft was at the same level as the host node. The HB portion primarily occupied the secondary brain. The PNC derived from the node graft was incorporated into the posterior host tissues. (e) and (f) The cases of posteriorly-positioned quail node grafts. Primarily, tissue that developed from the node graft was PNC, to which a host-derived narrow strip of SOX2-expressing neural tissue aligned. (C) An enlarged dorsal view of the area indicated in (A)(b). The anterior limit of the secondary brain leveled with the posterior FB of the primary brain. The dorsal part of the primary brain from the posterior FB to the MB (double arrow) was missing (arrowhead). (D) Immuno-fluorescence image of a section of the embryo at level “D” shown in (A)(d) superimposed on the phase-contrasted image. The grafted quail node developed into the floor plate (FP), PNC, and adjacent definitive endoderm (DE). The bars indicate 500 *μ*m in (C)(D).

Transplanting the quail nodes in the anterior epiblast field resulted in the secondary brain development underlain by the node-derived AME [Fig. 6(B)(a) to (c), AMEs shown in magenta]. However, they did not develop much of the PNC tissues [Fig. 6(B)(a) and (c), asterisks]. By contrast, the nodes grafted in the posterior epiblast field developed into the secondary PNC without producing AME [Fig. 6(B)(d)(e)]. These observations indicated that the secondary brain tissues are formed only when a node is grafted in the embryo position anterior to the node, where the AME develops.

Following the node grafts in the anterior epiblast field, the brain portions that developed in the secondary brains depended on the AP level of the graft position, changing from FB, MB to HB [Fig. 6B(a) to (c)]. This observation suggested that the regionally specified head precursors [Fig. 4(C)] developed into the brain tissues when the grafted node-derived secondary AME was supplied, reflecting the given regionalities. The embryos in Figure 6 represented cases in which the nodes were grafted at positions relatively distant from the embryo axis, resulting in some portions of the secondary brain developing separately from the primary brain. Grafting a node closer to the midline resulted in the fusion of two brain tissues at all AP levels, as shown in Figure 7 below, but the second brain regionalities were determined in the same way.

**Figure 7.**
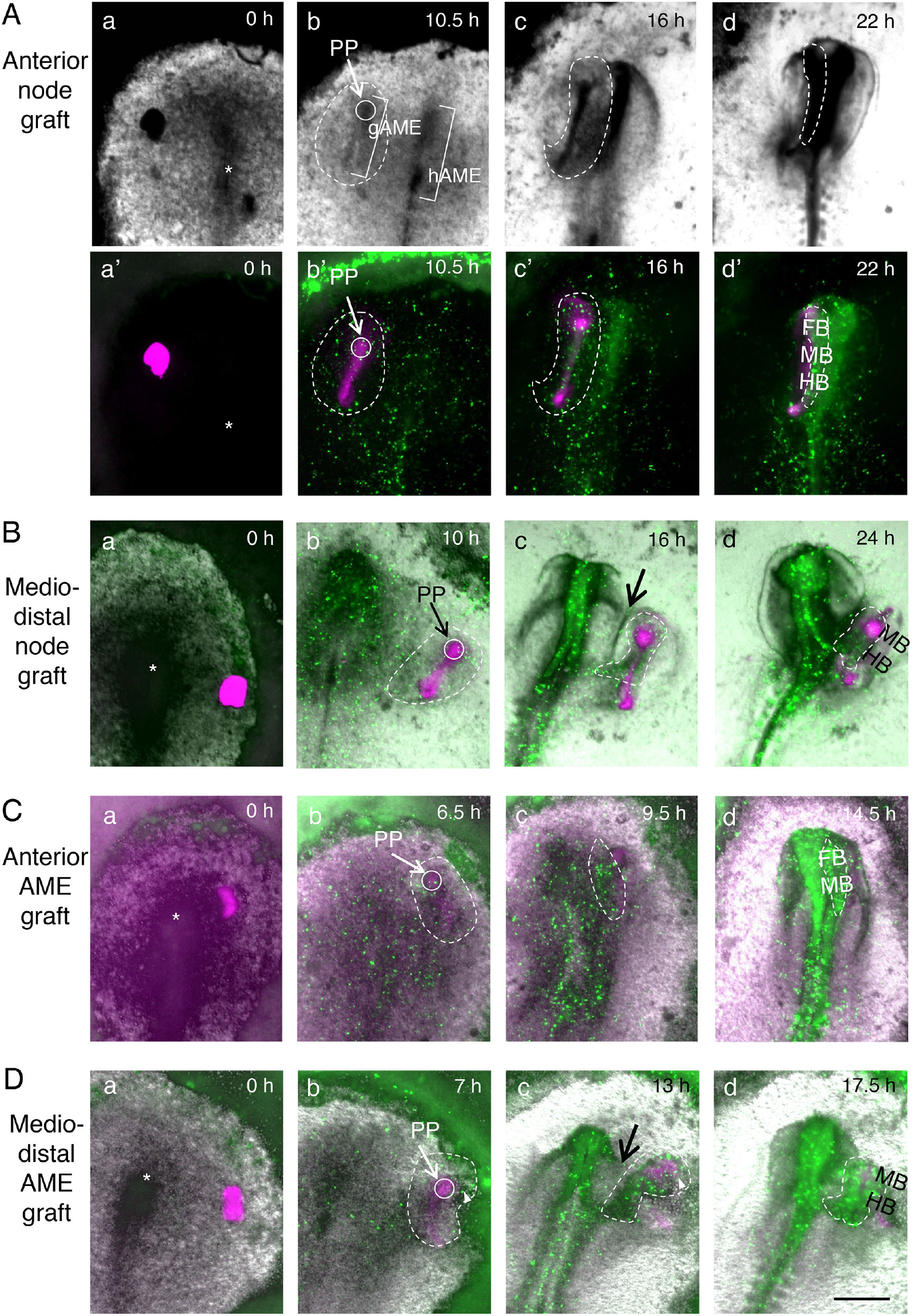
Live imaging of the events after grafting mCherry-transgenic quail node/AME in SN-EGFP-labeled st. 4 chicken embryos. The anterior is at the top. (A) Grafting a st. 4 Japanese quail node at an anterior level. (B) Grafting a quail node at a distal position of the host node AP level. (C) Grafting a st. 5 quail AME at an anterior level, similar to (A). (D) Grafting a st. 5 quail AME at a distal position of the host node level, similar to (B). In (A), upper bright-field images show changes in cell density (darker with a higher density) and lower images are composites of mCherry (magenta) and EGFP (green) fluorescence. In (B) to (D), bright-field, and fluorescence images are superimposed. All panels across (A) to (D), (a) to (d) show snapshots after h of culturing as indicated above. Panels (a) and (a’). The node/AME graft (magenta) and its position in the host embryo. The asterisks, the position of host nodes. Panels (b)(b’) in (A)(B). The stage when the node graft-derived AME [gAME with a bracket in (A)(b)] fully elongated to form prechordal plates (PP, encircled with an arrow) and more posteriorly positioned thinner anterior notochord. The gAME elongation occurred synchronously with host AME [hAME with a bracket in (A)(b)]. Panels (b) in (C)(D). The stage when the grafted st. 5 AME elongated and developed into PP and anterior notochord. Panels (b)(b’) across (A) to (D). The AME elongation coincided with the start of the convergence of surrounding head precursors (encircled by broken lines). Panels (c)(c’) across (A) to (D). The developmental stages when head precursors that converged to gAME started to form the secondary brain (encircled by broken lines). The frames were chosen for their representation of the event, rather than fixed time points. Panels (d) across (A) to (D). Later developmental stages for the node/AME-grafted embryos. The secondary brains (encircled by broken lines) fuse to the primary brains at all AP levels [(A)(C)] or their posterior end [(B)(D)]. When head ectoderm developed between the primary and secondary brains (arrows in (B)(c) and (D)(c)), the two brains remain separated. Arrowheads in (D)(b) and (c) indicate the cell originated from the area opaca contributing to the secondary brain. FB, forebrain; MB, midbrain; HB, hindbrain. The bar indicates 500 *μ*m.

When the secondary brain developed into FB to MB portions, they caused defects in the primary brain of the same portions (4/4), exemplified by Figure 6(B)(a); both primary and secondary FB portions lacked the facing dorsal tissues (open arrowheads). Figure 6(C) shows an enlarged dorsal image of the embryo in Figure 6(B)(b). Judging by the morphology, the secondary brain consisted of the posterior half of FB and full MB. Remarkably, the primary brain had defects in the dorsal brain tissues at the levels of secondary brain development. In the case of the specimen shown in Figure 6(B)(c) where the secondary MB developed without association with the primary brain, the MB portion of the primary brain was extraordinarily thin (open arrowhead). The observations of these and analogous embryos suggested that secondary FB and MB developed at the expense of precursors for the primary brain portions.

The graft level in the embryo shown in Figure 6(B)(d) was just an intermediate; the AME development from the node elicited secondary HB development, whereas posteriorly extended secondary notochord was incorporated into the host tissue. A cross-section of this embryo shown in Figure 4(D) indicated that the grafted quail node developed into the posterior notochord (PNC), floor plate (FP), and definitive endoderm (DE), consistent with earlier observations (Catala et al., 1996; Teillet et al., 1998) that FP, PNC, and abutting DE originate from the same precursor group. In this and analogous embryos having two PNCs, the neural plate widened and failed to close at the dorsal end. Grafting nodes in the posterior epiblast field resulted in graft-derived PNC development in isolation from the host spinal cord, as shown in Figure 6(B)(e)(d). In these cases, a host-derived narrow strip of neural tissues was associated with the graft-derived PNCs. We did not characterize them further but speculated that they were also spinal cord floor plates.

The graft-position dependent development and interaction of the grafted nodes with host head precursors were specific to st. 4 nodes, and not observed using st. 5 nodes. The grafted st. 5 nodes primarily self-differentiated into the PNC and somites (Supplementary Figure 4). This observation underscores the importance of the AME derived from st. 4 nodes in the production of secondary brain tissue.

### Live imaging of grafted nodes and AMEs for their development and interaction with head precursors

To investigate the sequence of events elicited by grafting nodes in the anterior epiblast field and resulting in the secondary brain development, we grafted st. 4 nodes from mCherry-expressing transgenic quail (Huss et al., 2015) into SN-EGFP-labeled chicken host (N = 13). This combination of fluorescence-labeled tissues allowed simultaneous imaging of the development of node graft-derived tissues and host cell migration under the influence of the graft.

A case of grafting a quail node in an anterior position of a host embryo is shown in Figure 7(A), whose full record is shown in Movie G1. In this figure, snapshots of bright-field images indicating changes in cell density (darker representing a higher density in the bright-field image), and combined fluorescence images are displayed in different panels. Panels (a) and (a’) indicate the initial state of grafting a transgenic quail node at a host position more anterior than the host node (asterisk). The grafted node developed in an elongated gAME synchronously with the host AME (hAME), which further differentiated into the anteriorly positioned PP and the thinner ANP in 10.5 h [Fig. 7(A)(b)(b’)]. Around this stage, the gAME-proximal host head precursors started to converge around gAME [encircled by white broken lines in Fig. 7(A)(b)(b’)], whereas those in the remaining region did so around hAME [Fig. 7(A)(b)(b’)]. These two groups of head precursors individually developed into brain tissues [a broken line encircles the gAME-centered group in Figure 7(A)(c)(c’)]. These developing brain tissues approached each other with time, reflecting the bi-centered convergence of the overall head precursors [Fig. 7(A)(c)(c’)]. The gAME-centered host epiblast cells developed into a thin secondary brain tissue covering FB to HB [Fig. 7(A)(c)(c’)], and eventually fused to the primary brain tissue that developed around hAME [Fig. 7(A)(d)(d’)]. In other embryos where gAME elongated shorter than this embryo, and the secondary brain contained only limited brain portions, for example, MB and HB, the fusion to the primary brain tissues occurred at these levels.

Figure 7(B) represents the case in which a quail node was grafted at an AP level similar to the host node, yet at a remote position (Movie G2). The bright field and fluorescence images were superimposed. The gAME elongated fully around 10 h of node grafting, when surrounding head precursors started to converge [encircled by a broken white line in Figure 7(B)(b)], similar to Figure 7(A). While these gAME-centered cells developed into the secondary brain tissue (encircled by a broken white line), the head ectoderm developed at the periphery of two cell convergences (thick arrow) and separated the primary and secondary brain tissues [Fig. 7(B)(c)]; the secondary brain tissue developed into independent MB/HB portions, and fused to the primary brain only at the posterior end [Fig. 7(B)(d)]. Examination of analogous cases indicated that the timing and extent of head ectoderm development between the primary and secondary brain tissues determine whether and how they fuse.

The above observations indicated that the gAME rather than the grafted node provides the center for head precursor cell convergence leading to the development of a secondary brain. To corroborate this model, we grafted AMEs of st. 5 transgenic quail embryos at host embryo positions analogous to Figure 7(A) and (B) [Fig. 7(C)(a) and (D)(a)], and performed live imaging [Fig. 7(C) and (D), Movies G3 and G4] (N = 12). The gAMEs elongated at the grafted position quickly to form PP and ANC in 7 h of grafting [(Fig. 7(C)(b) and (D)(b)), namely, 3 h earlier than the node graft-derived gAME [Fig. 7(A)(b) and (B)(b)]. The gAME-surrounding head precursors converged on gAME [Fig. 7(C)(b) and (D)(b)] resulting in secondary brain development [Fig. 7(C)(c) and (D)(c)], ending in the fusion of the primary and secondary brain portions [Fig. 7(C)(d) and (D)(d)]. Thus, grafting the st. 4 nodes or st. 5 AMEs elicited identical processes to form secondary brain tissues, with the exception of ~3 h earlier progression of the processes after st. 5 AME grafts. This 3 h difference is accounted for by the period required for the AME extension from the grafted node. This observation also indicates that the head precursors have a wide temporal range in the responsiveness to and convergence around a proximal AME.

To confirm the collective convergence of head precursors on gAMEs, we performed cell tracking analysis around two axes: the gAME axis (Graft axis) and the virtual axis (Vgraft axis) placed at a mirror-image position relative to the primary head axis (Supplementary Figure 5). The changes in the positioning of initially gAME-proximal cells relative to the Graft axis were compared with those of cells in the vicinity of the Vgraft axis, which provides a non-convergent control. Analyses of two embryo specimens under different contexts of gAME placement are shown. In the first embryo [Supplementary Figure 5(A)], an mCherry-transgenic quail node was grafted on a lateral side of the SN-EGFP-labeled chicken host, resulting in the secondary HB development. In the second embryo [Supplementary Figure 5(B)], a st. 5 quail AME was grafted at an anterior position of the host, resulting in the secondary FB and MB development. The tracking of SN-EGFP-labeled cells around the Graft axis during the cell convergence phase [7 h 20 min to 11 h 40 min for (A), and 3 h 10 min to 5 h 40 min for (B)] confirmed gAME-centered cell convergence, in contrast to the contralateral cell trajectories disregarding the Vgraft axis [Supplementary Figure 5(A’) and (B’)].

We also investigated whether the head precursor cells converging on gAMEs change their cell proliferation rate (Supplementary Figure 6). The epiblast cells of node-grafted embryos were labeled by 5-Ethynyl-2′-deoxyuridine (EdU) during 6 to 8 h, or 8 to 10 h after the graft. The EdU-labeled fraction of nuclei was compared among the following three regions: a region on the gAME, a region on the hAME, and a contralateral side region of gAME without an effect of AMEs. The labeling rates were equivalent among the regions in an embryo, 55.4% for 6–8 h post-graft labeling, and 49.8% for 8–10 h labeling, only showing a developmental stage-dependent decrease in the fraction of S-phase nuclei. Therefore, AME-centered cell convergence did not significantly change cell proliferation.

### Brain portions that developed in the secondary brains reflect the head precursor regionalities at the AME/node graft position

The observations reported above indicated that the grafted AMEs or node graft-derived AMEs direct surrounding head precursors to converge and allow them to develop into the secondary brains. Moreover, the graft positions of AME/node affected the resultant secondary brain portions, judging by morphological criteria. To ascertain the assignment of the secondary brain portions, we performed in situ hybridization analysis of embryos after 16–20 h of AME/node grafting at various positions in host embryos [Fig. 8(A)] for the expression of *Otx2* (for FB and MB) and *Gbx2* (for HB) [Fig. 8(B)]. Among *Otx2*-expressing tissues, the FB development was distinguished from MB by the lateral bulging. Four representative embryos are shown in Figure 8(B) with assigned second brain portions. The results obtained using in situ hybridization-based head portion determination were consistent with morphological brain portion assignments as shown in Figure 6.

**Figure 8.**
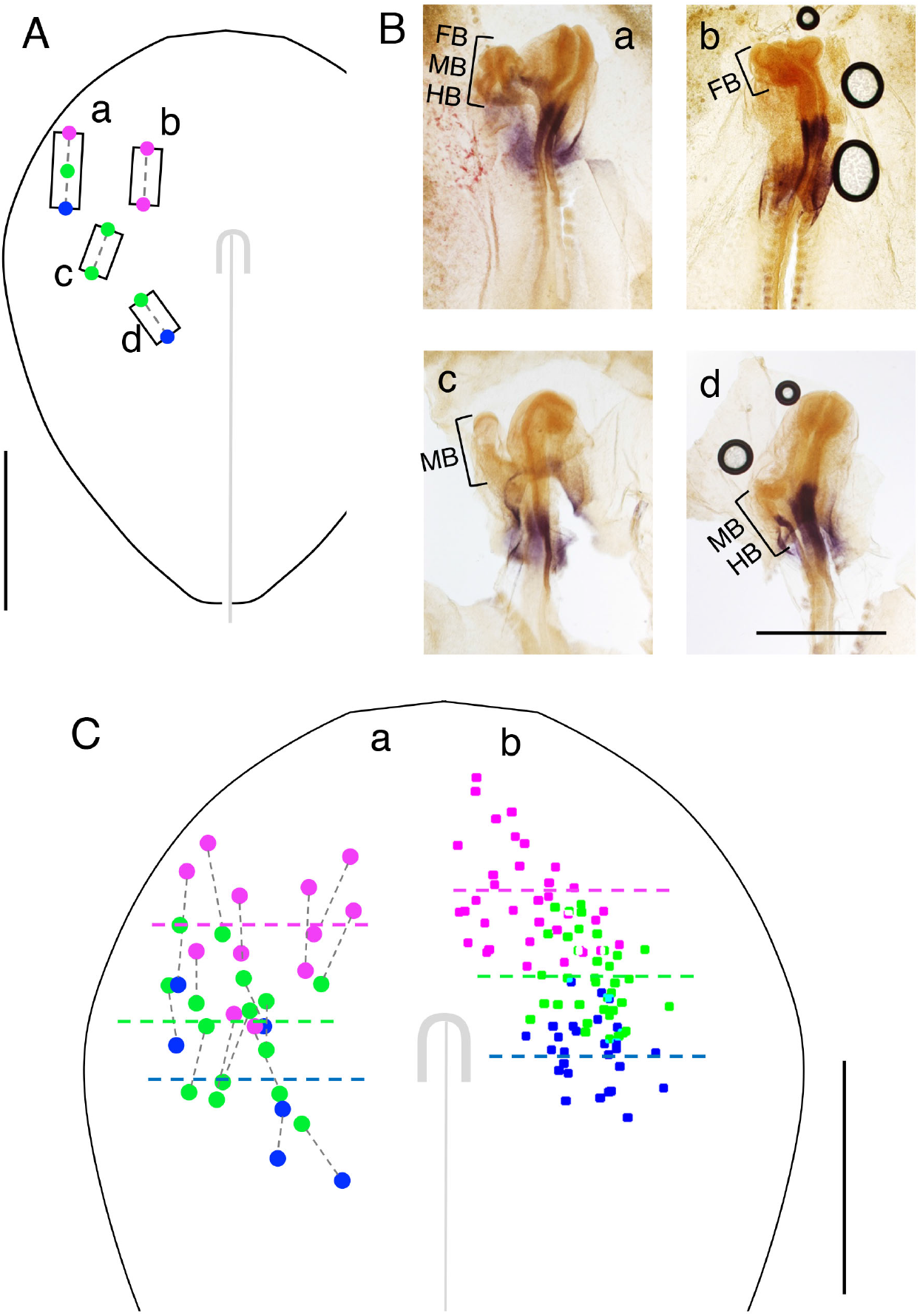
The gAME position-dependent development of the secondary brain portions. (A) Four examples of the gAME positions in st. 4 chicken host embryos showing graft positions and sizes shown on the left side. Graft (a) was a long strip of AME derived from an embryo of advanced st. 5. (B) After culturing for 16-20 h, the embryos were analyzed by in situ hybridization for *Otx2* (orange) and *Gbx2* (purple) to determine the development of FB, MB, and HB portions in the secondary brain. (a) The long gAME elicited secondary brain development containing all FB, MB, and HB portions. (b) The AME grafted more medially than (a) caused secondary FB development only. (c) This median level gAME caused only secondary MB development. (d) The posteriorly grafted gAME elicited secondary MB and HB. The secondary brain portions were color-coded for FB (magenta), MB (green), and HB (blue) and plotted on the anterior and posterior ends of gAME, except that gAME in (a) had additional midpoint MB data. (C) Comparison of the gAME position-dependent variations in the secondary brain portions with the head precursor regionalities shown in Fig. 4(C). (a) A summary of the color-coded secondary brain portions using data of 11 AME grafts and 5 node grafts. For the node grafts, we hypothesized 300 *μ*m anterior elongation of gAME from the grafted node and plotted the tissue codes on the hypothetical gAMEs. (b) The regionalities in the distribution of dorsal brain/proximal head ectoderm precursors shown in Fig. 2. Color-coded broken lines indicate average AP levels for individual plot groups. The anterior is at the top in all panels. The bars indicate 1 mm.

Then the assessed secondary brain portions were marked onto the anterior and posterior end positions of AME/node grafts at st. 4, in the way shown in Figure 8(A) [all those in panel (B) were AME grafts]. Next, we plotted the results using 16 gAMEs (12 AME grafts and 4 node grafts) in Figure 8(C)(a) (left panel), concerning the observed brain portions as determined by in situ hybridization in correlation with the gAME’s anterior and posterior end positions. Confirming the notion conveyed in Figure 6, the secondary brain portions that developed along with gAMEs strongly depended on the graft positions: Changing AME graft positions from anterior to more posterior positions resulted in the development of more posterior brain portions [Fig. 8(A)(B)(b)–(d)].

We compared this plot in Fig. 8(C)(a) with the head precursor regionality map [Fig. 4(C)] redisplayed in Figure 8(C)(b). The brain portion attributes were arranged similarly in the order and the distribution of AP groups in Figure 8(C)(a) and (b). The plots in (a) are positioned ~170 μm posterior to those in (b), as estimated by the AP level differences in the group averages. However, this difference is accounted for the post-grafting anterior elongation of gAMEs, which was not considered when drawing Figure 8(C)(a). The head ectoderm precursors are expected to be distributed in a broader lateral range than the those of the proximal head ectoderm [Fig. 8(C)(b)], as discussed in Figure 4(C). The wider lateral distribution of the plots in Figure 8(C)(a) than in (b) corroborates this notion.

Overall, the data in Figures 6–8 indicated that the graft position-dependent secondary brain development [Fig. 8C(a)], and the prospective AP levels of head tissues after convergence on the head axis in normal development [Fig. 8C(b)] are strongly correlated. As shown in Figure 7, the effects of gAMEs involve first directing the surrounding head precursors to converge on gAME, and then allowing them to develop into the secondary brain portions. Figure 8 shows that the secondary brain portions develop according to the traits of AP regionality of the head precursors, which in normal development is manifested primarily by the contribution to the head ectoderm of the same AP group as the node-proximal brain precursors [Fig. 4(A)]. This observation supports the following model: (1) The head precursors consisting of the anterior epiblast are bipotential concerning their development into either brain tissues or covering head ectoderm. (2) Once the AME tissue develops in the vicinity of head precursors, they converge centering on the AME, where centrally positioned precursors develop into neural tissues and those at more distal positions into the head ectoderm. (3) Regardless of which tissue they develop into, they possess gross regionalities to orient themselves to which AP levels of tissues they contribute. (4) However, the positioning in a cell group at a critical developmental stage determines the details concerning the tissue portions to which they eventually contribute, a corollary of the results shown in Figure 3.

## DISCUSSION

### Basic patterns of collective migration of epiblast cells visualized by SN-EGFP-labeled epiblast

We conducted this study because of our interest in clarifying the discrepancy between the broad anterior epiblast region of *Sox2* N2 enhancer activation suggesting neural potency (Iwafuchi-Doi et al., 2011; Uchikawa et al., 2003) and the reported brain precursor region, which is much narrower and confined to the proximity of the node (Fernández-Garre et al., 2002).

Previous anterior epiblast fate analyses (e.g., Fernández-Garre et al., 2002; Schoenwolf and Alvarez, 1991) employed homotopic grafting of labeled tissue plugs and analyzed the distribution of the plug-derived tissues the following day. We felt that live imaging analysis at single-cell resolution is required to accurately describe the cellular events giving rise to the head tissues, i.e., the brain and head ectoderm, with the AP differentiation into FB, MB, and HB levels. We thus developed a SN-based technique to label epiblast cells randomly in a scattered fashion with EGFP.

Figure 2 shows three primary analyses based on tracking of SN-EGFP-labeled epiblast cells: (A) drawing trajectories, (B) characterizing developmental stage-dependent changes in the direction and rate of epiblast cell migration at different embryo positions, and (C) determining the time-dependent deformation of the epiblast cell sheet. This deformation includes the compression/extension vertical to the AP axis, extension along the AP axis, and changes in cell density, which is inversely correlated with the surface area.

The trajectories of SN-EGFP-labeled cells [Fig. 2(A)] indicated a couple of essential features of the epiblast cell migration. (1) First, the cell migration patterns are distinct between the pre-node and post-node AP levels. Anterior to the node, the initial cell migration was directed primarily anteromedially (toward the embryo axis), followed by the medial-directed convergence. The cells initially located at the periphery of the epiblast field migrated over a distance of 1 mm toward the axis. A more detailed analysis of cell trajectories allowed the drawing of maps of the head tissue origins of different AP levels (Figs. 3 and 4), as discussed below. As the cell migration in the anterior epiblast is minimal between st. 4 and st. 5 (Supplementary Figure 1), maps that are drawn for the st. 5 anterior epiblast can be regarded as equivalent to st. 4 maps. (2) Second, immediately posterior to the node level, many cells converged on the embryo axis, and then changed direction toward the posterior end, resulting in the massive posterior flow of cells along the posterior midline [Fig. 2(A) and (B)]. This group of cells constituted the posterior domain of the N2-enhancer-active region of epiblast [Fig. 4(D) and Movie E]. Because our primary interest was in brain development, these cells were not investigated further. However, the posterior flow of N2-enhancer-positive epiblast cells has an essential bearing on the trunk-level neurogenesis, which will be discussed below. (3) Despite observing these basic features, the details of cell migration patterns were not stereotyped, but showed a significant level of variation between the embryos [Fig. 2(A)(B)] and even between the sides of an embryo [Fig. 2(B)(C)]. The last feature reflected a considerable level of flexibility in the epiblast cells concerning their developmental fates, which will be discussed below.

### The flexibility of anterior epiblast cells concerning brain or head ectodermal fate, confirmed by the supply of gAME as the second cell convergence center

The analyses in Figure 2 indicated that the most extensive cell migration in the anterior epiblast field, converging on the extending node-derived AME, occurs between st. 5 and st. 6, a period of ~4 h, which formed the anterior neural plate. Following the trajectories of individual cells in the presumptive brain portions at st. 6 back to st. 5 successfully identified the original cell positions. The cells for future brain portions were arranged in the same AP order in the epiblast field and were proximal to the brain axis extending anteriorly from the node (Fig. 3B). However, the distribution of these cells was highly variable among the embryos and between the embryo sides. The case of a highly asymmetric distribution of brain precursors shown in Figure 3(B)(b) accounts for an interesting observation reported by Fernández-Garre et al. (2002) that an epiblast graft at one side of a st. 4 embryo occasionally contributes to both sides of the brain tissues. Analysis of cell migration between st. 5 and st. 6 indicated that the incidental positioning at st. 6 (when the neural plate is formed) determines the cell’s fate to develop into the brain or head ectoderm, and into which brain portions or head ectoderm of equivalent AP levels. Averaging data for four such embryos and different sides, the map of brain portion precursors at st. 5 matched well with the earlier data obtained using transplanting tissue plugs (Fernández-Garre et al., 2002; Schoenwolf and Alvarez, 1991).

The above observation strongly suggested that the brain/head ectoderm dichotomy is highly context-dependent. Indeed, the precursors for later-stage (st. 8–9) dorsal brain and proximal head ectoderm were similarly mapped into zones for each brain portion at st. 5 [Figs. 3(C), 4(C)]. These zones merged with the st. 6 brain precursor distributions and extended anterolaterally. The overall head precursor distributions occupied most of the N2 enhancer-active anterior epiblast field [Fig. 4(D)].

To determine whether all of these dorsal brain and head ectoderm precursors have comparable developmental potentials as the node-proximal brain precursors, we introduced the second gAME of Japanese quail origin to the particular region of the anterior epiblast. The second AME was supplied by grafting either a st. 4 node, which extends gAME in the same way as the host AME [Figs. 6, 7(A)(B)] or an AME isolated from a st. 5 quail embryo [Figs. 7(C)(D), 8]. In the former case, the choice of an embryo precisely at st. 4 as the node donor was essential, as detailed in Figure 5.

After the supply of gAME, regardless of by grafting a st. 4 node or st. 5 AME, the host epiblast cells converged on the gAME to produce a thicker epiblast zone compared with the surroundings (Fig. 7). This process was similar to the cell sheet thickening observed around the hAME [Figs. 2(C), 7]. Then, the secondary brain tissues developed in the central region of the thickened cell sheet. This observation demonstrates that the broad anterior epiblast has the potential to develop into either brain tissue or head ectoderm. The observation also indicates that the head tissues develop from the anterior epiblast in two steps. The first step is to produce an AME-centered thick tissue via cell convergence. In the second step, the cells centrally located in the convergent cell population develop into brain tissues, whereas more peripherally located cells develop into the head ectoderm [Figs. 2(C), 7].

The *Sox2* expression patterns at various developmental stages (Uchikawa et al., 2011) indicated that strong *Sox2* expression that occurs broadly in the anterior epiblast at. st. 5 was still maintained in the head ectoderm region at st. 7. This observation may suggest that the anterior epiblast-derived cells maintain the brain vs. head ectoderm bipotentiality to a certain degree even after tissue territories are established. Only later do the cells in the head ectoderm territories downregulate *Sox2*. In addition, in mouse embryos, a fraction of the cells that initially express *Sox2* downregulates it and develop into head ectoderm (Cajal et al., 2012). In the chicken embryo brain region, the D1 enhancer of the *Sox2* gene is activated at st. 6 confined to the neural plate (Iida et al., 2020). This enhancer activity is maintained in the later neural tube and neural crest. The D1 enhancer activation may direct the cell fate to brain tissues, while N2 enhancer activation provides the brain potential.

It was interesting to determine whether known diffusible factors, particularly those emanating from the AME, are involved in these two steps of head development. As AMEs secrete antagonists against various signaling molecules, we grafted COS7 cells overexpressing LEFTY1, CERL1 (both Nodal antagonists), DKK1 (Wnt antagonist), or NOGGIN (BMP antagonist) underneath the anterior epiblast. However, none of their combinations mimicked AMEs in eliciting cell convergence or promoting the development of brain tissue (data not shown), only adding to the list of factors ineffective for promoting neural development as reported by Linker and Stern (Linker and Stern, 2004). Pinho et al. (2011) reported a putative RNA binding protein gene *Obelix* expressed in the cell convergence phase of the epiblast. Analysis of the regulation of this gene may clarify the mechanisms in the AME-directed epiblast convergence.

### AP regionalities in the epiblast field and determination of the brain portions

Grafting a node or node-derived AME thus elicited convergence of the epiblast toward the gAME and secondary brain development. It was remarkable that the brain portions at the secondary site reflect the regionality of the graft site in the epiblast field that was determined by tracing back the head portion levels at st. 8 to 9 (Figs. 6 to 8). For instance, when a gAME was placed in the epiblast region with the FB and MB regionalities, brain tissue consisting of FB and MB portions but lacking HB developed. These observations indicated that the epiblast cells that converged on gAME develop into brain portions only according to their intrinsic regionalities. In other words, the AME primarily acts as the epiblast converging center and does not contribute much to the AP specification of the convergent epiblast.

However, these regionalities are likely macroscopic, and the contribution of individual epiblast cells into which portions or sub-portions may be determined by cell position-dependent interaction with surrounding cells. This mechanism was suggested by the embryo-dependent significant variations in the distribution of st. 5 precursors giving rise to the st. 6 brain portions (Fig. 3).

In the case of the MB and HB boundary, it has been shown that initially grossly set brain regions are finely tuned by mutually repressive interactions between the transcription factors OTX2 and GBX2 (Katahira et al., 2000; Martinez-Barbera et al., 2001). Analogous processes will take charge of organizing the macroscopic tissue regionalities into the precise brain tissue architecture.

### The continuity of the anterior epiblast into the anterior proximal region of area opaca

Examination of the trajectories of SN-EGFP-labeled epiblast cells in many embryos indicated that the cells travel freely across the boundaries between the anterior area pellucida (thought to develop into the embryo proper) and the proximal region of the area opaca (thought to develop into extra-embryonic tissues). The regions of the cell tracks indicated by broken ovals in Figure 2(A)(a) show such examples. These observations suggested that the proximal region in the anterior area opaca can be regarded as an extension of the embryogenic epiblast. Indeed, we observed cases where the cells originating from an area opaca region contributed to the brain tissues that developed around the gAME [e.g., Fig. 7(D) arrowheads].

Many earlier node graft experiments were performed, selecting the proximal regions of the area opaca (Storey et al., 1992) or germinal vesicles at the epiblast anterior end as grafting sites (Dias and Schoenwolf, 1990). These experimental maneuvers assumed that these regions are not specified, and hence bear no embryo regionalities. The development of brain portions after the node grafts has been interpreted on this assumption.

However, Streit et al. (1997) reported the following interesting observation concerning the responsiveness of the area opaca. The region of the area opaca that produced host-derived brain tissues was limited to the region with L5 antigen expression, extending anteriorly from the area pellucida [Fig. 9(A)]. In addition, the capacity of the area opaca to produce neural tissues was lost after st. 5, concomitant with the loss of L5 expression. The region of the area opaca with L5 expression appears to overlap with the region where SN-EGFP-labeled cells migrate seamlessly from and to the area pellucida [Fig. 2A(a) and Movies A, B1, E, G2 to G4].

**Figure 9.**
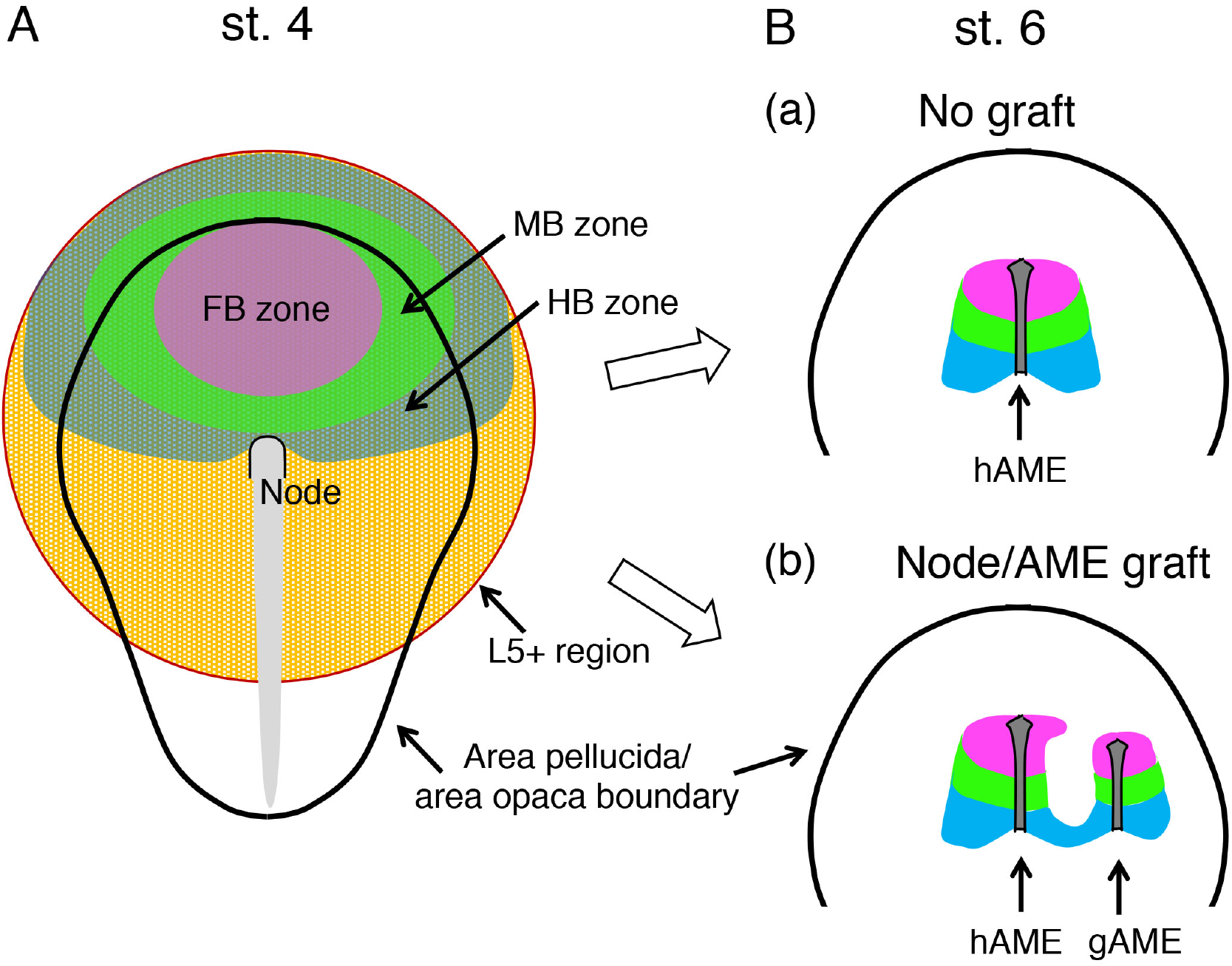
The model of anterior epiblast regionalities with the potential to develop into FB, MB, and HB portions or abutting head ectoderm of the equivalent AP levels. (A) Distribution of the head tissue developmental potential of anterior epiblast at st. 4. The head tissue developmental potential extends over the boundary of area pellucida to the L5+ domain of area opaca (Streit et al., 1997). The zones for the developmental potential for brain portions are drawn using pale colors, based on the data shown in Figs. 4 and 7, and data reported in earlier studies (Streit et al., 1997). (B) The convergence phase of the anterior epiblast. (a) The case without exogenous gAME supply. The epiblast cells lateral (in the sense of the AP level) to the endogenous node-derived AME (hAME) converge on the hAME, maintaining the original regionalities, indicated by the full-color codes. (b) The case with node graft-derived gAME or directly grafted gAME. In these cases, two AMEs, hAME and gAME, provide two convergent centers for the epiblast cells. The epiblast cells between the two AMEs are split into two converging groups toward the two different AMEs, resulting in the deficits in the head precursors at the facing sides of the forming head. Presumably, because the ventral brain portions develop from the epiblast population abutting the AME, the precursor deficits result in the lack of dorsal brain portions and the head ectoderm, as observed in Fig. 5(A)(a) and (C).

Considering the fact that grafting the node in the L5-expressing area opaca region resulted in the development of MB and HB tissues, this region must be given the MB and HB regionalities. Incorporating this and the above points and extending the data in Fig. 4, we propose a head tissue regionality map of st. 4 epiblasts, as shown in Figure 9(A). This model also accounts for the brain portions that developed following the st. 4 node grafts onto the anterior margin of the area pellucida or proximal zone of the area opaca (Dias and Schoenwolf, 1990; Storey et al., 1992). Knoetgen et al. (Knoetgen et al., 2000) reported that grafting the nodes from the mouse or rabbit embryos at the area pellucida/area opaca boundary of st. 4 chicken embryo resulted in brain tissue development similar to the avian node grafts. The use of *Otx2* expression as a FB/MB indicator by these authors clearly showed the AP orientations of these secondary brain tissues. The prominent features were that the anterior ends of the secondary brain tissues always pointed to the host forebrain and that the *Otx2*-expressing secondary forebrain always developed in the area pellucida, in contrast to the *Gbx2*-marked hindbrain that developed in the area opaca, fully supporting the model shown in Figure 9(A). Data reported by Pinho et al. (2011) confirmed that the epiblast convergence-dependent cell thickening accompanying the *Obelix* expression occurred in response to the chicken node graft close to the area pellucida boundary.

### Competition for head precursors between two AMEs that act as the cell convergence centers

An important message from this study is that an essential function of an AME is to provide the convergence center for the epiblast lateral to the AME [Fig. 9(B)(a)]. This model predicts that if a gAME is provided at a position lateral to hAME, then the epiblast cells between two AMEs are split into two groups converging into the two different AMEs [Fig. 9(B)(b)]. The live imaging of SN-EGFP-labeled epiblast cells under such conditions supports the split in the direction of cell migration (Fig. 7). In other words, hAME and gAME compete for the pool of converging epiblast in the small region between the two AMEs.

Observation of the defects in the primary brain structure associated with the secondary brain development [Fig. 6(B)(a)–(c)] likely indicated the outcome of the competition of cell sources between the AMEs. The lack of the dorsal brain portions and head ectoderm shown in Figure 6(B)(a) and (C)(b) suggests the following process. Presumably, because the ventral brain portions develop from the epiblast population abutting the AME, the precursor deficits caused the defects in the dorsal brain and head ectoderm portions. As the regionalities of the epiblast cells are maintained during these processes [Fig. 9(B)(b)], the dorsal tissue defects occur at specific brain portion levels, as most clearly shown in Figure 6(C).

### Implications from the massive posterior cell migration along the midline at the post-node level

One remarkable observation in Figure 2(A), though not analyzed in-depth in this paper, was the massive convergence of cells on the midline at the post-node level, and their subsequent posterior migration along the embryo axis (broken rectangles). These cells originated from the cells in the posteriormost zone with N2 enhancer activity occupying the ~200 μm AP thickness and overlapping with the HB precursor distribution, as shown in Fig. 4(D) and Movie E. The N2-enhancer-positive cells primarily contributed to the brain portions (Uchikawa et al., 2003), yet a significant contribution to the spinal cord was also observed (Iwafuchi-Doi et al., 2011; Kondoh et al., 2016). These observations suggest that a fraction of such cells that once activated the *Sox2* N2 enhancer contribute to the posterior axial structure as either neural progenitors or NMPs. In a clonal analysis study of post-gastrulation mouse embryos, Tzouanacou et al. (2009) observed several clones giving rise to the brain tissues and posterior neural/paraxial tissues with a vacancy at the intermediate axial levels. The clones of this pattern could be accounted for by the rapid posterior cell migration along the embryo axis of a subclone having a brain subclone counterpart. This process is under investigation, but the cell trajectory data in Figure 2(A) alone suggest that the cell population that first converges on the embryo axis and then migrates posteriorly makes an essential contribution to the trunk development.

## CONCLUSIONS

In the first part of this study, we developed a technique to randomly label epiblast cells by SN-EGFP to trace these cells in head development by live imaging. The single-cell-resolution analysis of cell trajectories and brain precursor positioning in the epiblast field provided a new fate map of brain portions. The analysis suggested considerable flexibility of epiblast cells concerning their contribution to the brain or head ectoderm.

We thus performed further analysis on the anterior epiblast regionality using node or AME grafts. The combination of SN-EGFP-labeled epiblast and mCherry-expressing node/AME was adopted to perform live imaging of the epiblast–graft interactions. The overall picture given by the second analysis was as follows: The anterior epiblast cells are bipotential for the development into brain tissues or head ectoderm but carry gross regionalities corresponding to the brain portion levels. These epiblast cells converge on the nearest AME, regardless of the hAME or gAME. Among the epiblast cells that have converged, those centrally positioned develop into brain portions according to their regionalities. In contrast, others develop into the head ectoderm of corresponding regionalities. These results renew the regulatory mechanisms underlying head tissue development after st. 4, and the roles of the node in this process.

In a recent review, (Martinez Arias and Steventon, 2018) carefully analyzed the widely considered “organizers” in various animal species for their activities. They concluded that the classical amphibian organizer is “a contingent collection of elements, each performing a specific function,” which are dispersed temporally and spatially in embryos of other species. Our observations on the avian nodes support their proposal concerning the need for reconsideration of the organizer concept.

## ACKNOWLEDGEMENTS

We thank Rusty Lansford for his provision of the *mCherry*-transgenic Japanese quail line. This study was supported by JSPS KAKENHI Grant JP16H06280 (Advanced Bioimaging Support), Grants-in-Aid for Scientific Research JP26251024 and JP17H03688 to HK, National BioResource Project “Chicken/Quail,” and Private University Research Branding Project of the Ministry of Education, Culture, Sports, Science and Technology (MEXT) Japan, as well as by the Kyoto Sangyo University Research Fund for Institute for Protein Dynamics.

## COMPETING INTERESTS

We declare that no financial or non-financial competing interests exist.

## MATERIALS AND METHODS

### Fertilized eggs

Non-transgenic chicken and quail fertilized eggs were obtained from local breeders. Fertilized eggs of *mCherry*-transgenic Japanese quail (TG(PGK1:H2B-chFP) (Huss et al., 2015) were provided by the Avian Bioscience Research Center at Nagoya University as part of the National Bioresources Program. The embryos were staged according to (Hamburger and Hamilton, 1951) and (Ainsworth et al., 2010). Animal experimentation was performed in accordance with the guidelines of Nagoya University and Kyoto Sangyo University.

### Random EGFP labeling of the epiblast by electroporation of Supernova vector cocktails

A st. 4 chicken embryo was isolated using a ring-shaped filter paper support attached to the vitelline membrane, and placed between a pair of 2 × 2 mm electrodes of 6 mm distance in the orientation of the ventral side upward facing the anode, using a set-up described previously (Kondoh and Uchikawa, 2008; Uchikawa et al., 2003; Uchikawa et al., 2017). The following mixture of vector DNAs was prepared [Fig. 1(A)]: 1 μg/μl pK031 (TRE-CRE-pA) (Mizuno et al., 2014), 0.5 μg/μl pK038 (CAG-LoxP-Stop-LoxP-GFP-IRES-tTA-pA) (Mizuno et al., 2014), and 0.4 μg/μl pTK-CAG-tTA (provided by Y. Takahashi), delivered at a volume of ~2 μl between the epiblast and vitelline membrane, and electroporated with four pulses of 8 V with 50 ms duration and 1 s intervals using CUY21SC (NEPAGENE, Ichikawa, Japan). The embryos were placed on the agar-albumen medium (Uchikawa et al., 2003) and cultured in the orientation of the ventral side upward. The embryo images were recorded with time-lapse imaging at 10 min intervals (except for Movies G1 and G2, at 25 min intervals) using a humidified chamber at 38°C under an AF6000 inverted microscope (Leica, Wetzlar, Germany).

Linear level adjustment of color channels, pseudo-color operations, and conversion of Tiff file stacks into Avi format time-lapse movies were performed using FIJI software (Schindelin et al., 2012). The movies were produced at a rate of 6 frames/s, showing 1 h of real time in 1 s.

### Node and AME grafting

To graft nodes, stage-matched chicken host and Japanese quail node donor embryos, both at st. 4, were isolated using the ring filter support and placed on agar-albumen support used in embryo culture. The donor node was excised as rectangular tissue block by an operation using a sharpened tungsten needle from the ventral side and transferred to the host embryo. A size-matched rectangular well was formed in the host embryo, into which the donor node was inserted [Supplementary Figure 2(A)(B)].

To isolate the AME from st. 5 Japanese quail embryo, a rectangular incision along the AME contour was made from the ventral side, leaving the epiblast intact. Then, 5 μl of 2.5% Pancreatin (Wako-Fujifilm, Japan) in HEPES-buffered saline (Qiu et al., 1998) was added to the incision and kept at room temperature for < 1 min to detach the AME from the epiblast. The stripped AME was rinsed and inserted between the epiblast and the ventral layer of a st. 4 chicken host embryo through an incision at an anterior aspect of grafting in the ventral layer [Supplementary Figure 2(C).

To record early cell convergence responses of the host epiblast, Supernova vector electroporation was performed at early st. 4 of the recipient embryo, and after 2 h of incubation when the host was still at st. 4, the above graft operations using an mCherry-transgenic donor were performed.

### Trajectory analyses of SN-EGFP labeled cells

The image sequences were first smoothened using a median filter (r = 1) and Gaussian blurring (sigma = 1.0). To manage the variation in fluorescence intensity between the frames and fluorescence image radius among the labeled cells, software implemented in C was developed in-house to perform the following procedures. (1) Local fluorescence maxima were collected as the candidate cell positions of labeled cells. (2) These cell positions were ordered according to their peak intensities. (3) Donut-shaped masks of 2 px width were prepared with 2 px radius increment steps spanning 4–20 px to collect intensity distribution data around the maxima. (4) These donut masks were applied to every peak to test the significance of the peak position intensity compared with peripheral regions using a ROKU algorithm (Kadota et al., 2003). The peaks that passed this test were registered as the “cells.” (5) The test was repeated by increasing the donut radius to collect all possible cells with various fluorescence diameters. (6) The information on the labeled cell positions was transformed using an R-script to align the horizontal axis of the forming head and keep the st. 5 node position at its posterior end. Then, trajectories of the SN-EGFP-labeled cells were drawn over image sequences using software in C developed in-house [e.g., Fig. 2(A)]. To draw cell trajectories relative to the gAME axis, the frames were reoriented accordingly. This protocol successfully tracked the labeled cells over tens of frames (covering several hours) but created gaps when the fluorescence signal of a cell became lower close to background noises in consecutive flames. When more extended cell tracking was needed, as shown in Figure 3(C), the tracks were generated by visually monitoring cell positions on the frames processed in the following way. The cell position data were generated to include low-fluorescence spots allowing the appearance of fluctuating background spots and overlain the original fluorescence image, allowing the choice of consistent tracks by ImageJ (Schneider et al., 2012) plugin developed in-house.

The optical flow that describes the local image feature displacements was computed by searching the coordinates for local intensity maxima that minimize the cross-correlation coefficient between immediate frames. Grids at constant intervals, corresponding to 150 or 250 μm in real space, were set so that a grid point corresponds to the posterior end of the head axis and points align with the axis. For each grid point, a mean optical flow was calculated by averaging vectors located within a circle of the radius equivalent of the grid intervals [e.g., Fig. 2(B)]. To illustrate the epiblast field’s sequential deformation, the new grid points in the following frame were calculated using the mean optical flow vectors in sequence. Deforming lattice representations of changing grid points were drawn using cubic spline interpolation of the lines connecting the original square lattice points [e.g., Fig. 2(C)].

### Whole-mount in situ hybridization of embryos

Embryos were fixed with 4% PFA in phosphate-buffered saline (PBS) at 4°C overnight and kept in methanol at –20°C. Whole-mount in situ hybridization was performed as described by (Henrique et al., 1995). The *Gbx2* and *Otx2* probes were synthesized using relevant templates (Katahira et al., 2000) (provided by H. Nakamura), labeled by digoxigenin (DIG) and fluorescein isothiocyanate (FITC), respectively, and used at 500 ng/ml. *Gbx2* hybridization was detected by purple color development following incubation with a 1/2000 dilution of alkaline phosphatase-conjugated anti-digoxigenin antibodies (Roche, Basel, Switzerland), washing, and incubation in 340 μg/ml nitroblue tetrazolium chloride and 175 μg/ml 5-bromo-4-chloro-3-indolyl-phosphate in 0.1 M Tris-HCl, pH 9.5, 0.1 M NaCl, 50 mM MgCl_2_, and 0.1% Tween 20. Subsequently, *Otx2* hybridization was detected by orange color development using a 1/500 dilution of alkaline phosphatase-conjugated anti-FITC antibodies (Roche) and HNPP Fluorescent Detection Set (Roche). After color development, embryos were mounted in RapiClear 1.52 (SUNJin Lab, Hsinchu, Taiwan) and photo recorded using an Axioplan 2 microscope (Zeiss, Jena, Germany).

### Whole-mount immunostaining of embryos

Chicken embryos of st. 8 to 10 were fixed by 4% paraformaldehyde (PFA) in PBS for 1 h at 4°C. The fixed embryos were stored in methanol at −20°C. The embryos were placed sequentially in 25% methanol, in 0.1% Tween 20 in PBS, and finally in 10% Blocking Reagent (Roche), 0.5% TritonX-100, and 0.1% Tween 20 in PBS, in which immunoreactions were performed. Primary antibodies were rabbit anti-SOX2 (Epitomics, Burlingame, CA, USA), used at 1/500) and mouse QCPN (quail cell marker antibody, Developmental Studies Hybridoma Bank, Iowa University, used at 1/100), and reacted at 4°C for 36 h (anti-SOX2) or 18 h (QCPN). After rinsing the embryos in 0.1% Tween 20 in PBS four times at room temperature, the secondary antibodies (Alexa 488/568-labeled donkey anti-rabbit/mouse IgG, Abcam, Cambridge, UK) were applied at 1/1000. The fluorescence images were taken using an M165 FC fluorescence stereomicroscope (Leica, Germany).

### 5-Ethynyl-2′-deoxyuridine (EdU) labeling of node-grafted embryos

The embryos isolated with a ring filter paper support and with a node graft at st. 4 were incubated at 38°C in the orientation of the ventral side upward. After 6 or 8 h, the embryos were flipped for the dorsal side facing upward, the vitelline membrane was pierced, 200 μl of 40 μM EdU was added to the epiblast side of the embryos, and the embryos with this orientation were incubated for further 2 h. Only the epiblast cells were labeled using this procedure. We confirmed that these dorsal-side-upward cultures allowed embryo development at least up to st. 15, similar to the ordinary ventral-side-upward cultures. After fixation with 4% paraformaldehyde for 15 min at room temperature, the embryo specimens were processed for Alexa488 fluorescence using the EdU Proliferation Assay Kit (Abcam) and counterstained with 1 μg/ml 4′,6-diamidino-2-phenylindole (DAPI). Fluorescence images were taken using an FV3000 inverted laser microscope (Olympus, Tokyo, Japan).

## SUPPLEMENTARY FIGURES

**Supplementary Figure 1.**
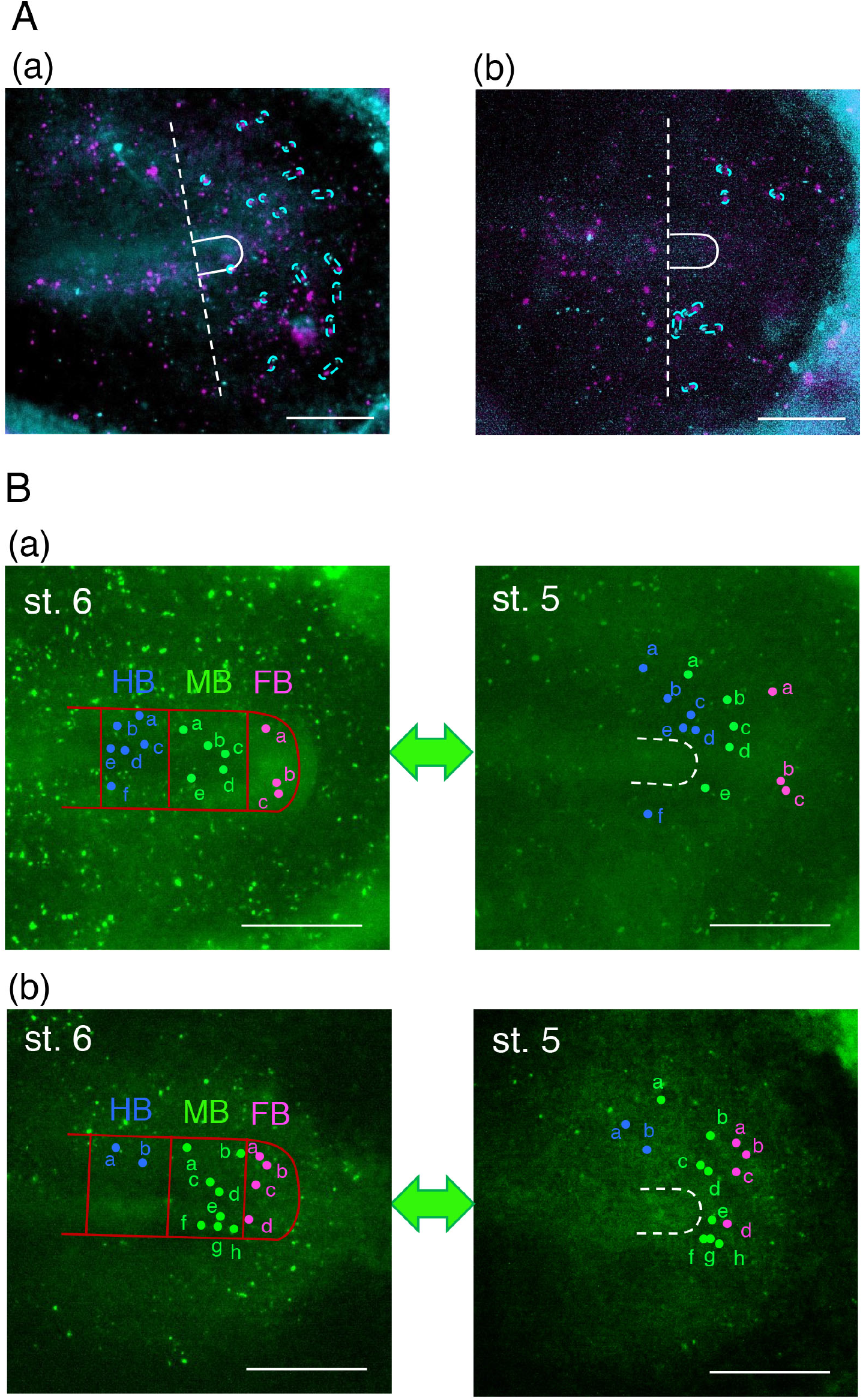
Additional cell tracking data between st. 4 and st. 6. (A) Comparison of the SN-EGFP-labeled cell positions between st. 4 and st. 5 in two representative embryos. The fluorescence images at st. 4 in cyan and those in st. 5 in magenta were superimposed. 24 conspicuous spot pairs representing drifts in the cell positions anterior to the node levels (broken lines) between the stages were chosen (encircled in magenta broken lines), and their displacement distances were measured, giving an average of 39 ± 26 *μ*m displacements in 130 min without defined orientations. The bars indicate 500 *μ*m. (B) The distribution of the precursors for individual brain portions of two additional embryos at st. 6 and st. 5. Annotations and color codes are the same as in Fig. 3(B). Embryo (a) was the same embryo as in Fig. 2(A). The embryo (b) was used in movie E, where the SN-EGFP labeling was more efficient on the left side (upper in the panel). The bars indicate 500 *μ*m.

**Supplementary Figure 2.**
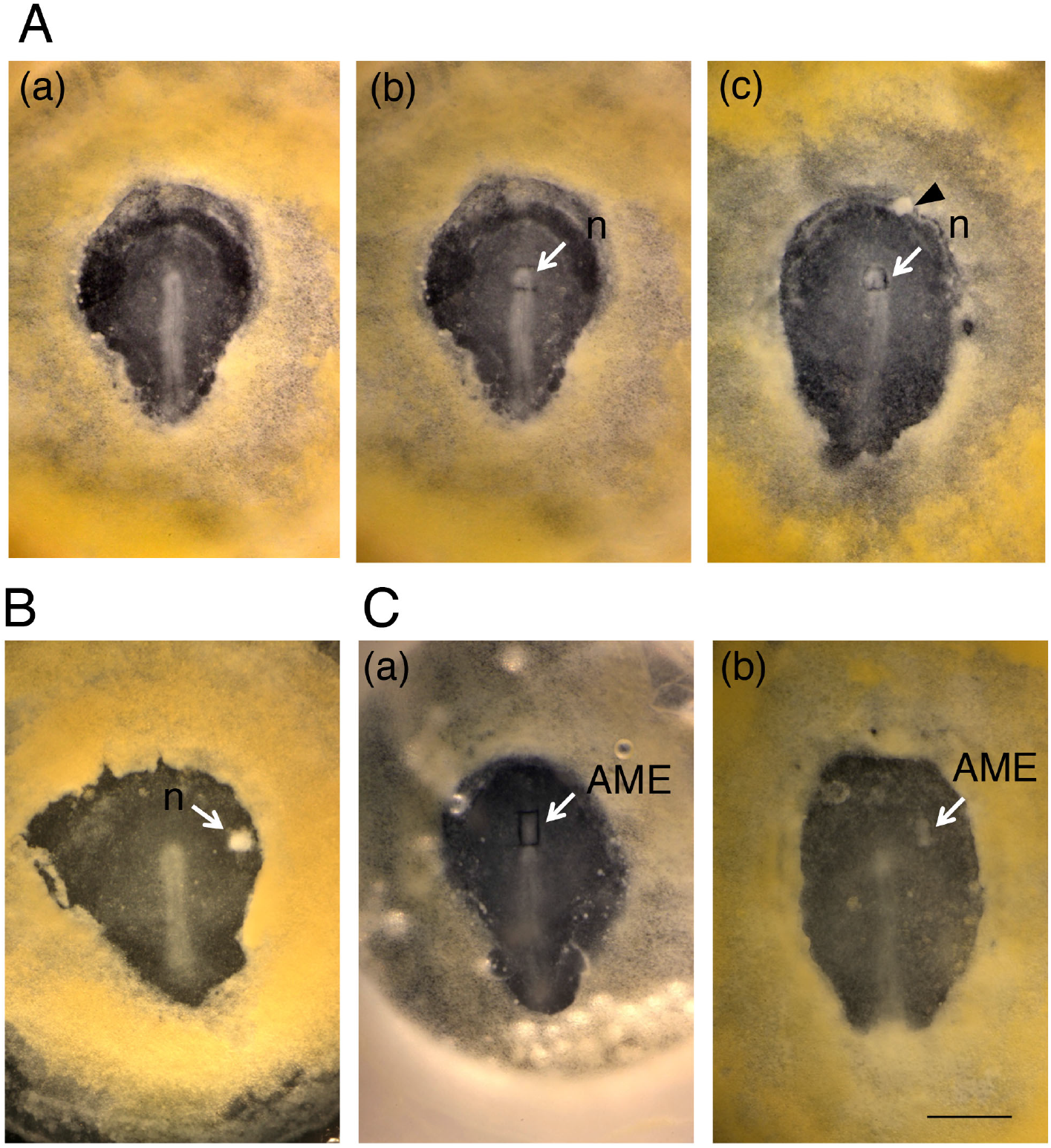
The procedure to graft a Japanese quail node/AME in a st. 4 chicken host embryo. (A) Homotypic node graft. (a) A Japanese quail embryo at st. 4, shown in Figure 5(A)(c) in the ventral view. (b) The node tissue (n with an arrow) was excised by making incisions at four sides. (c) The quail node (n with an arrow) was pushed into the square hole after node excision in the chicken host embryo. The excised host node to be removed is seen on the top left (arrowhead). (B) An example of a node graft in an epiblast field. The data of this embryo is shown in Figure 6(B)(b). (C) An example of grafting AME excised from a st. 5 donor Japanese quail embryo. (a) A rectangular incision made around the AME (arrow). (b) The AME-grafted host embryo. See Materials and Methods for details of the procedures. The data of this embryo is shown in Figures 7(B) and 8(B). The bar indicates 1 mm.

**Supplementary Figure 3.**
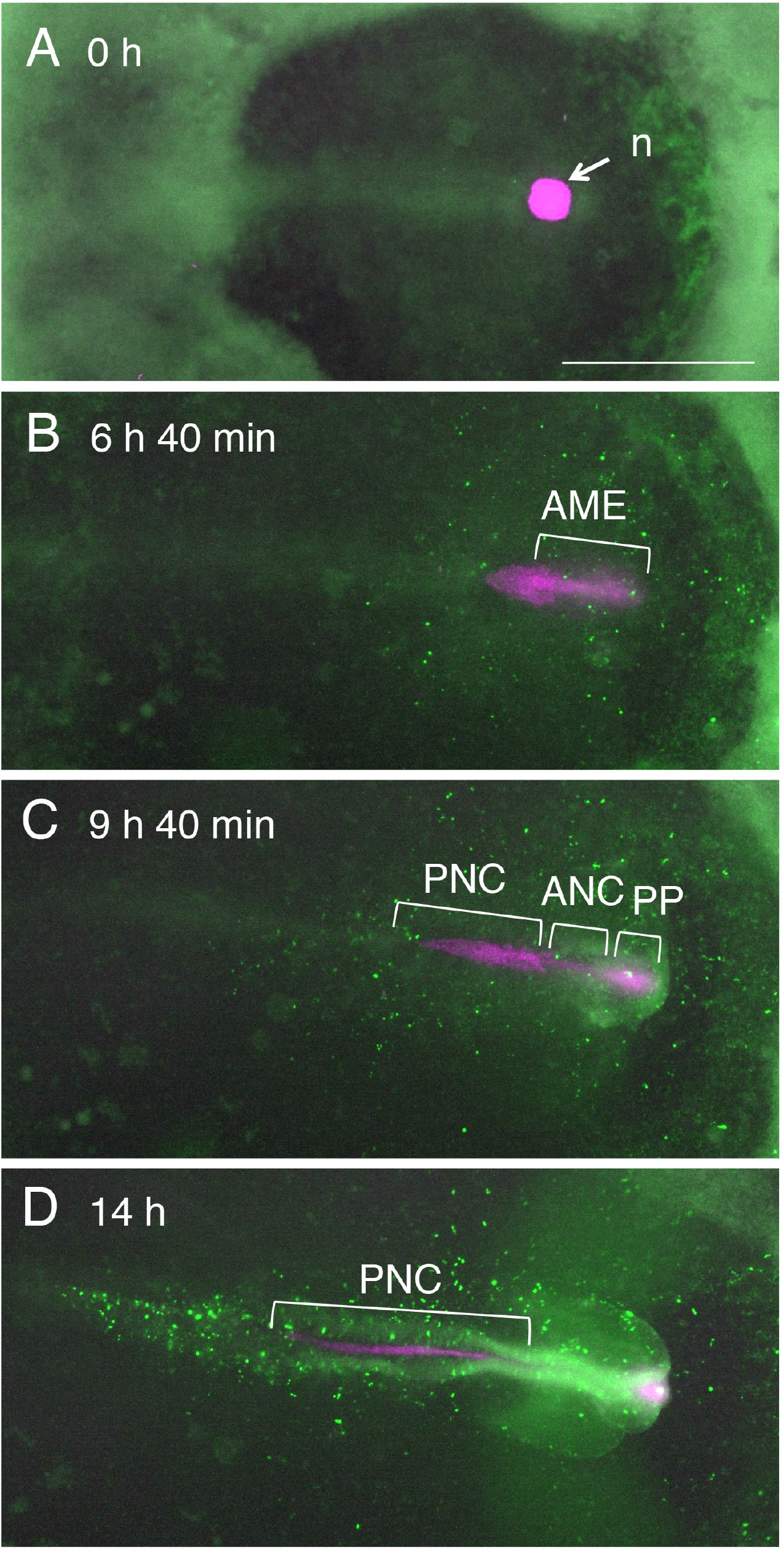
Development of mCherry-labeled Japanese quail node in homotopically grafted SN-EGFP-labeled chicken embryo. (A) Shortly after the node graft. mCherry fluorescence is shown in magenta. (B) 6 h 40 min after the graft. Anterior mesendoderm (AME) developed first. (C) 9 h 40 min after the graft. The AME developed into the prechordal plate (PP) and anterior notochord (ANC). Development of the posterior notochord (PNC) was underway. (D) 14 h after the graft. PNC elongated. The bar indicates 1 mm. All frames are shown in Movie F.

**Supplementary Figure 4.**
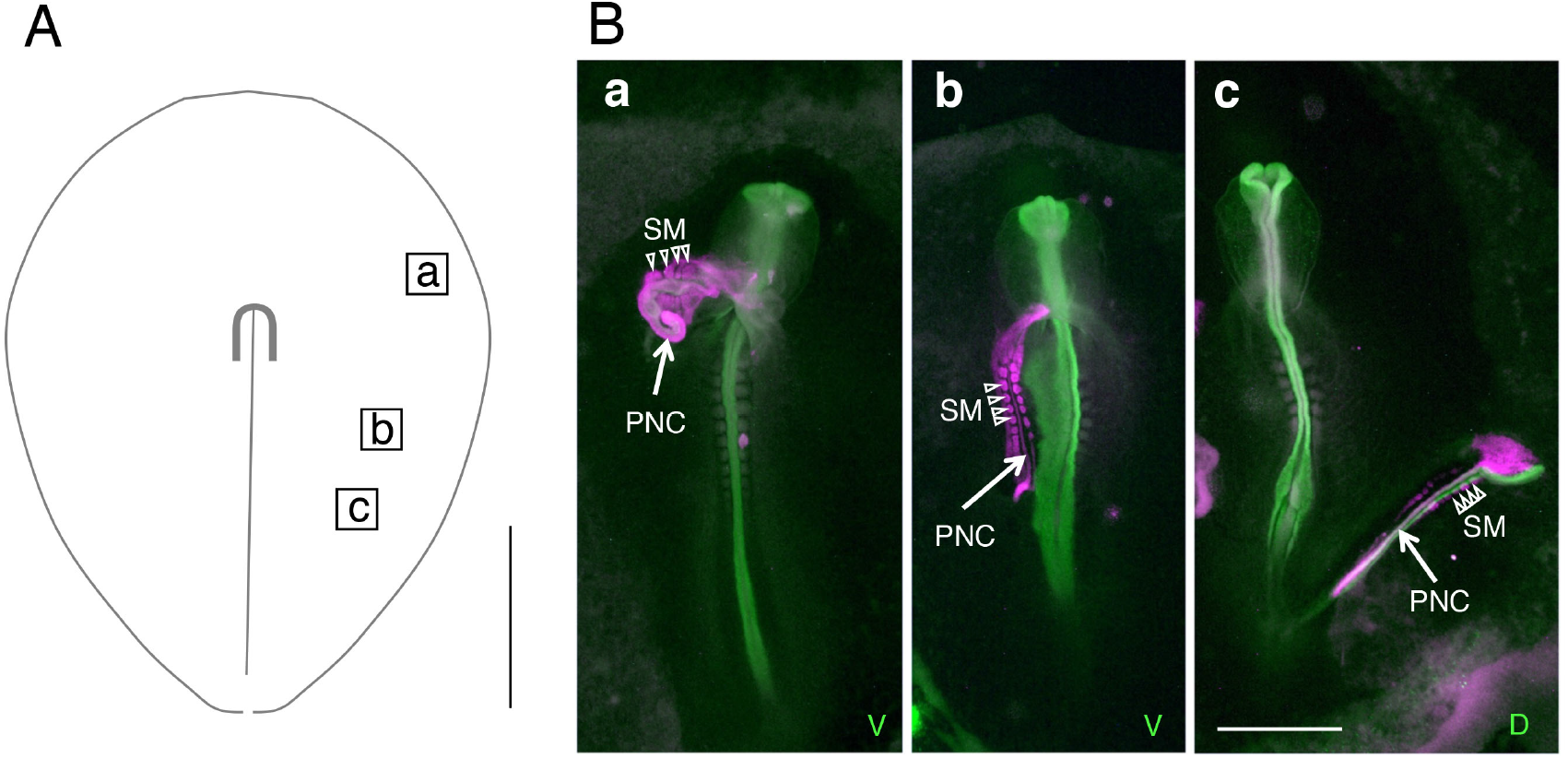
The outcome of st. 5 node grafts at various anteroposterior levels of st. 4 embryos. (A) Positions of the node grafts (a) to (c) (rectangles). (B) Embryos after culturing for 16-20 h, stained for SOX2 (green) and quail-derived tissues (magenta). In all embryos, the grafted node self-differentiated into the posterior notochord (PNC) and the somites (SM). (a) The tissues derived from the anteriorly positioned graft were not incorporated but protruded from the host tissue. (b) Medial position graft resulted in the fusion with the host tissue at the anterior end. (c) Grafting at the posterior-distal position resulted in node-derived posterior notochord and somites independent of the host tissue. A narrow strip of host-derived neural tissue associated with the secondary notochord, analogous to Fig. 6(B)(e)(f). The bars indicate 1 mm.

**Supplementary Figure 5.**
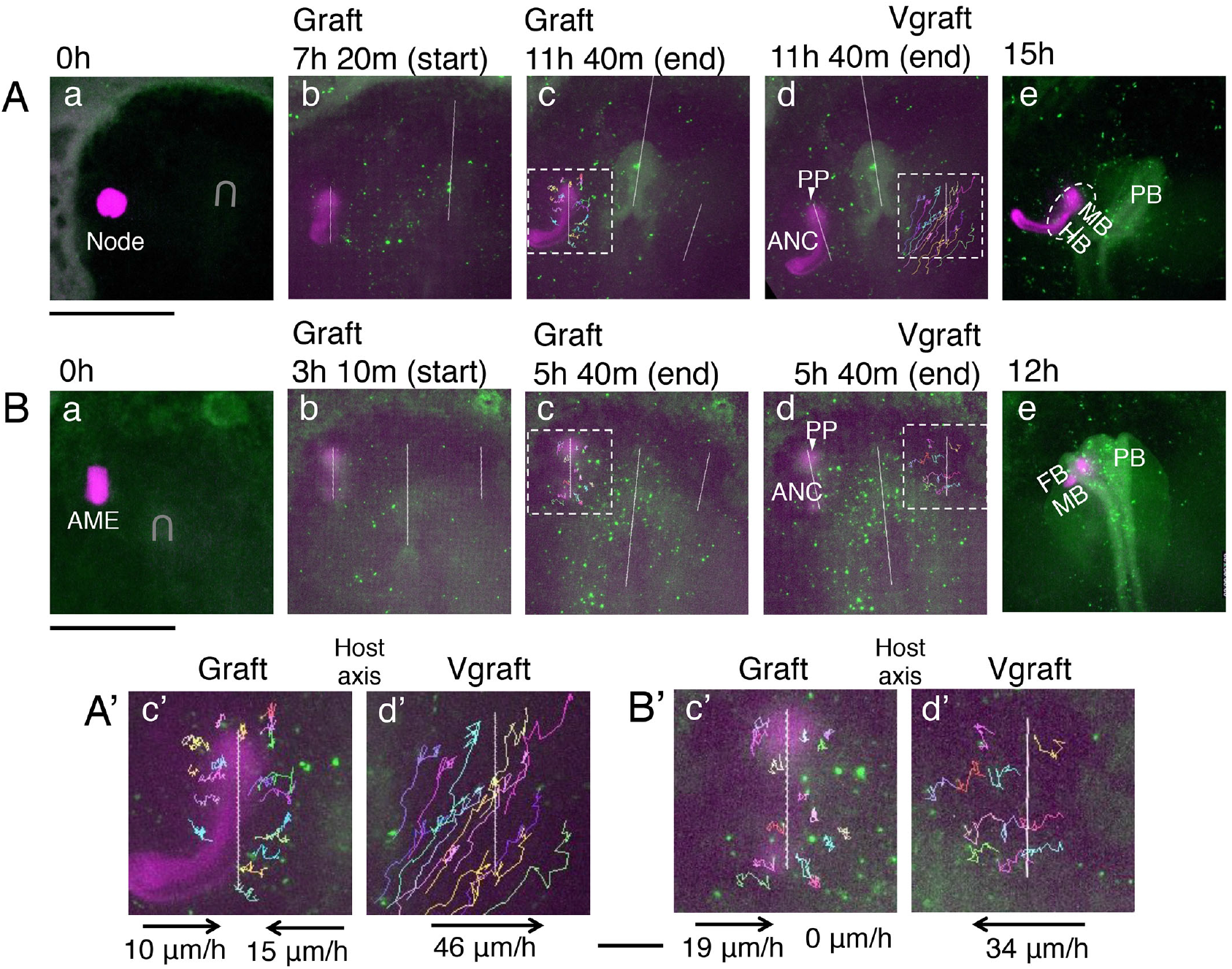
Confirmation of head precursor convergence on gAMEs by cell track analysis. (A) An SN-EGFP-labeled chicken embryo was grafted at early st. 4 with an mCherry-expressing quail node at a lateral position of the host node. The secondary MB and HB developed. (B) A chicken embryo grafted at late stage 4 with st. 5 quail AME at an anterior position. The secondary FB and MB developed, which eventually fused to the primary brain. Two axes were set on the epiblast plane: the gAME axis (Graft) and the virtual axis (Vgraft) placed at a mirror-image position relative to the primary brain axis. (a) Embryos shortly after the graft of a node (A) or an AME (B) indicated in magenta. U-shape indicates the host node position. (b) The initial time point of cell track measurement. The left line is the graft axis, which was kept vertical and at a fixed position during cell track analysis. (c) At the end of cell track analysis. The areas of cell tracks around the node/AME grafts enclosed by broken rectangles are enlarged in (A’)(c’) and (B’)(c’). (d) The cell tracks around the Vgraft axis. (e) Embryos at later stages of development. FB, MB, and HB indicate secondary brain portions in the proximity of the graft-derived prechordal plate (PP) and anterior notochord (ANC). PB, primary brain. (A’) and (B’). The details of the labeled cell tracks around the Graft axis and Vgraft axis. Average cell displacements relative to these axes are indicated by the arrows (direction) and the migration rate (μm/h). The bars in (A) and (B) indicate 1 mm. The bar for (A’) and (B’) indicates 200 μm.

**Supplementary Figure 6.**
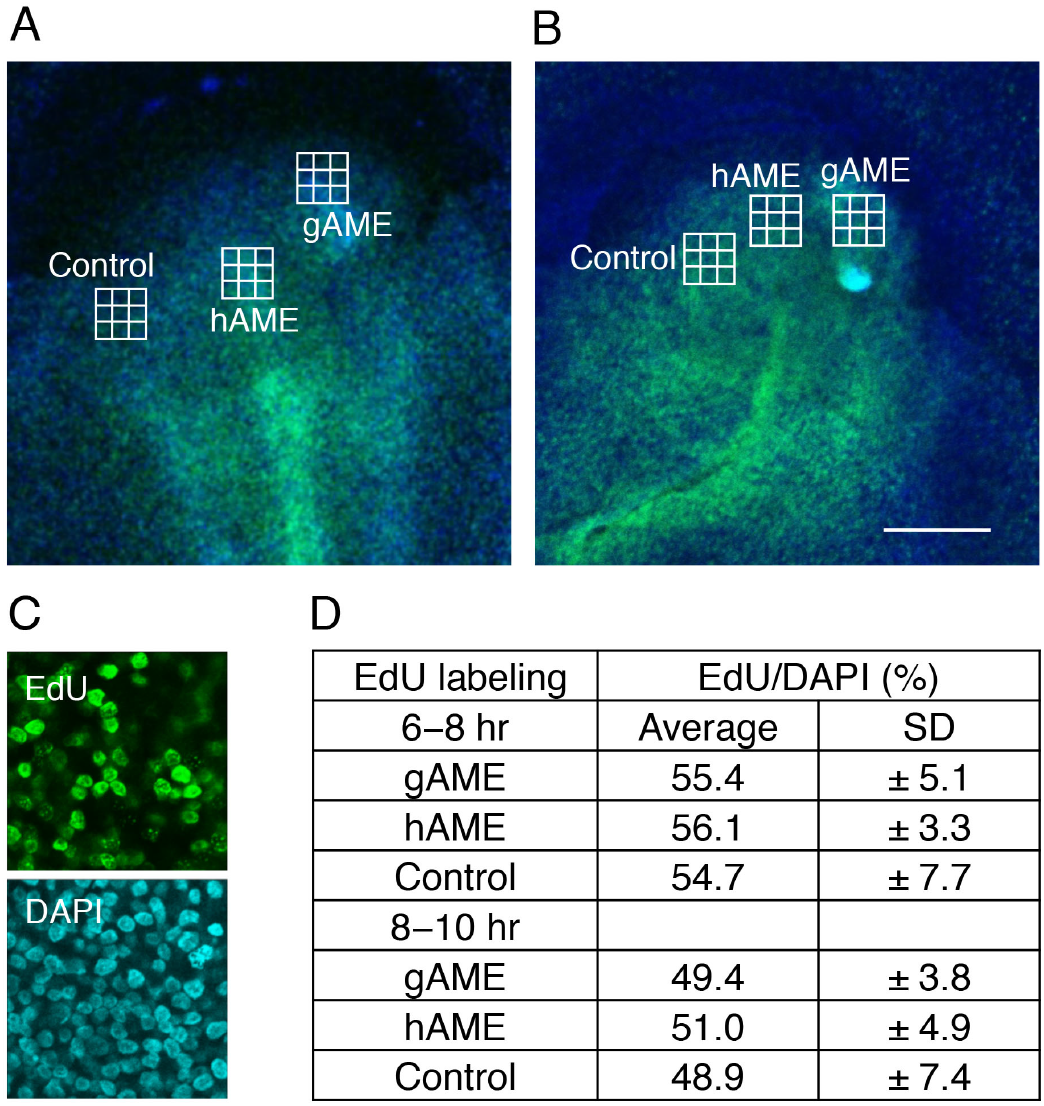
Detection of S-phase nuclei by EdU labeling of node-grafted epiblasts. (A) The epiblast of an embryo labeled with EdU after 6 to 8 h after grafting the node with DAPI nuclear staining. (B) An embryo labeled 8 to 10 h after grafting the node. After the EdU labeling, embryos were fixed and processed for Alexa 488 and DAPI fluorescence. Three rectangular regions of 234 × 234 μm^2^ covering the node graft-derived AME (gAME), host AME (hAME), and on the contralateral side of gAME (AME-unaffected control) were chosen for analysis in each embryo. Each region was divided into nine subregions to determine the fraction of EdU-labeled nuclei in all DAPI-stained nuclei. The bar indicates 500 μm. An example of such a subregion comparing EdU labeling and DAPI staining is shown in (C). An average of nine subregions in each measurement region is shown in (D) with standard deviations (Source data 2). An EdU labeling frequency of 56% was estimated for the embryo in (A) (6-8 h labeling) and 50% for (B) (8-10 h labeling).

## MOVIES LIST

Frame intervals were 10 min, except Movies G1 and G2, which were with 25 min intervals. The scale bars indicate 1 mm. Representative frames are shown as snapshots.

**Movie A.**
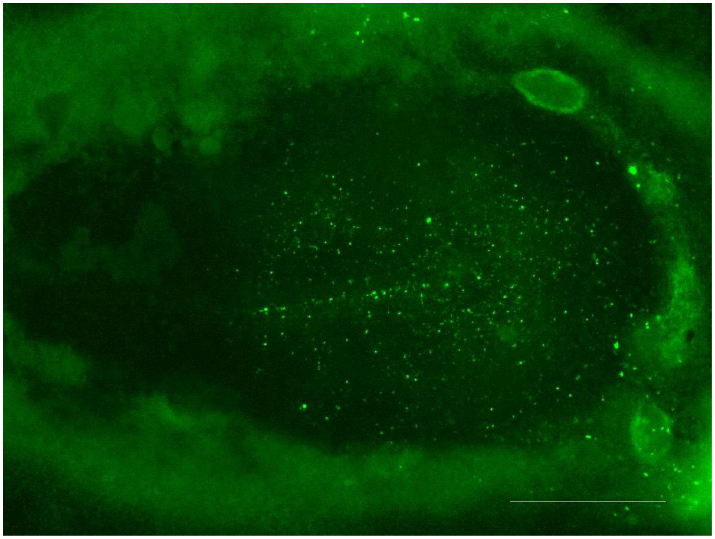
Original data of Figure 1(B). (Frame 24)

**Movies B1 and B2.**
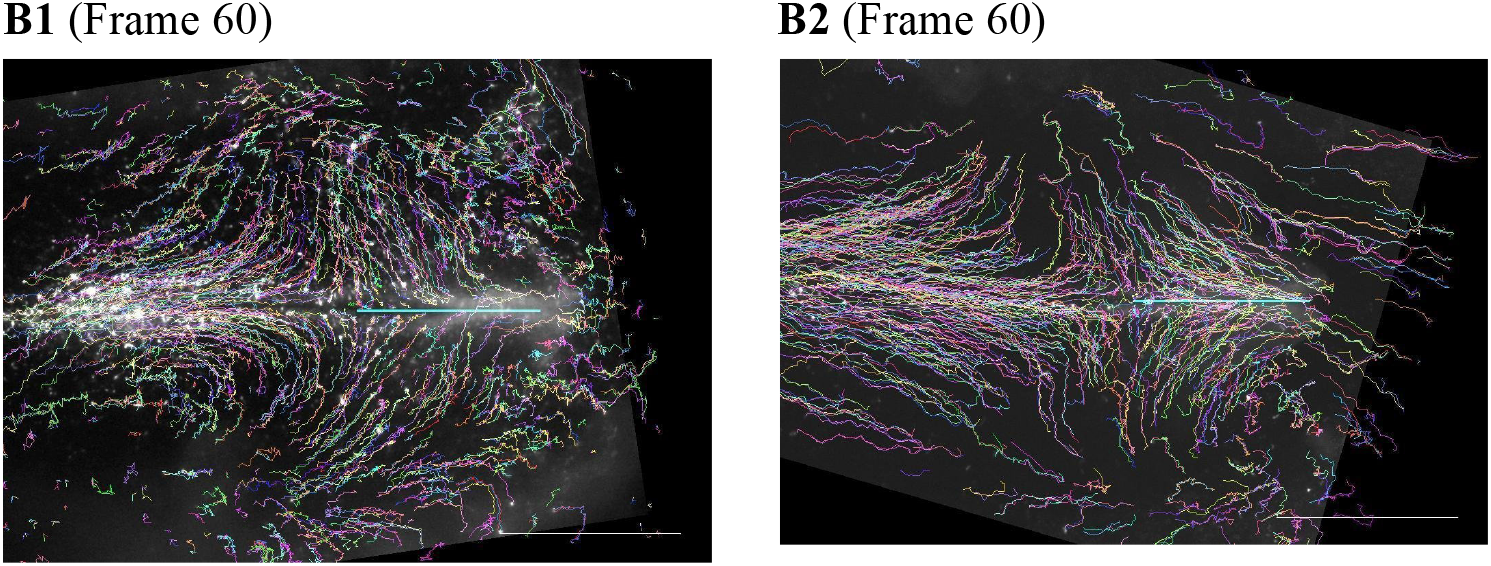
Tracking SN-EGFP-labeled cells in Fig. 2(A)(a) and (b).

**Movies C1 and C2.**
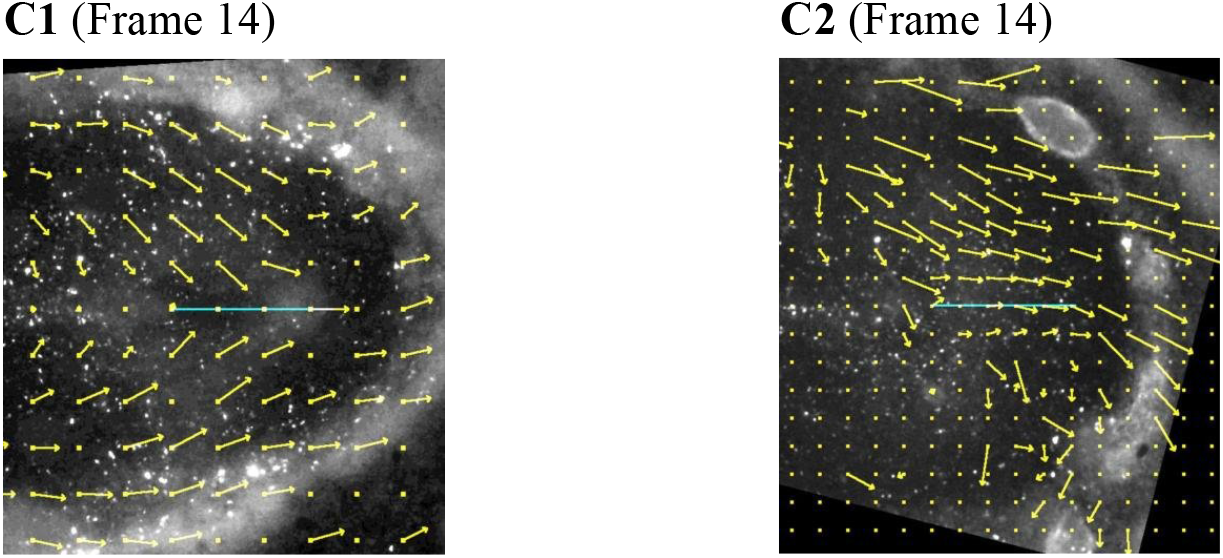
Cell migration vectors in Fig. 2(B)(a) and (b).

**Movies D1 and D2.**
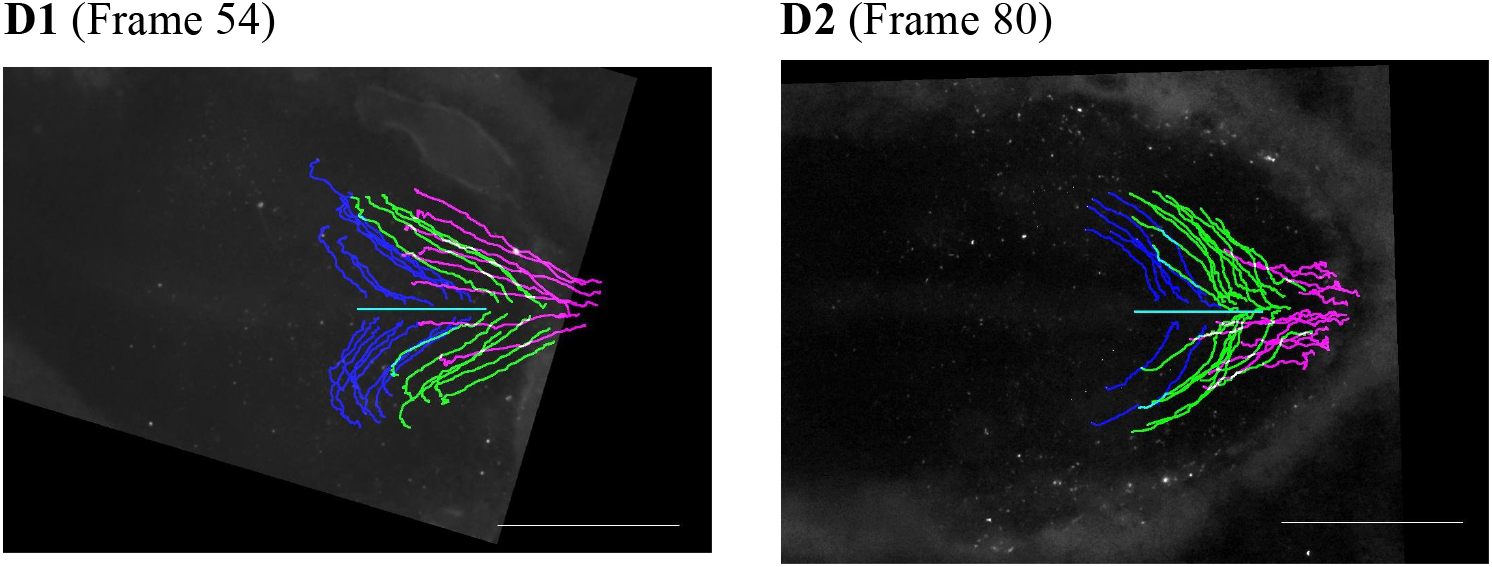
Backtracking data for Fig. 3(C) and the second example.

**Movie E.**
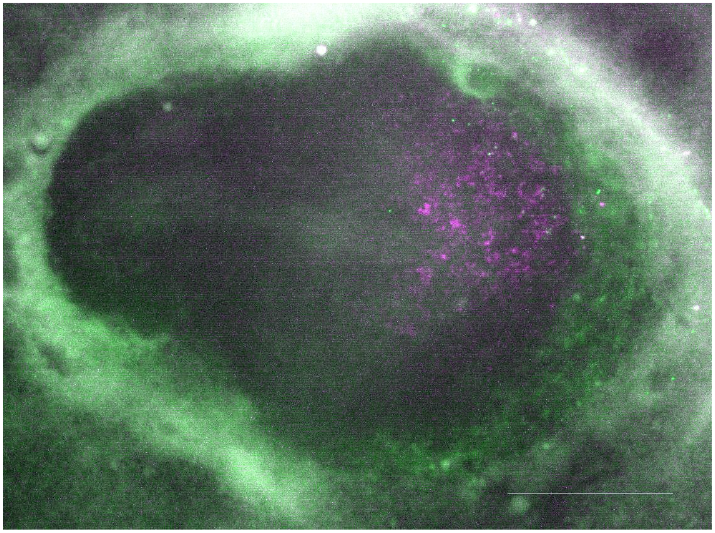
Comparison of N2 and SN-EGFP for Figure 4(D). (Frame 24). Vector DNA electroporation was focused on the embryo left side to show the N2 enhancer activity at the margin of anterior epiblast.

**Movie F.**
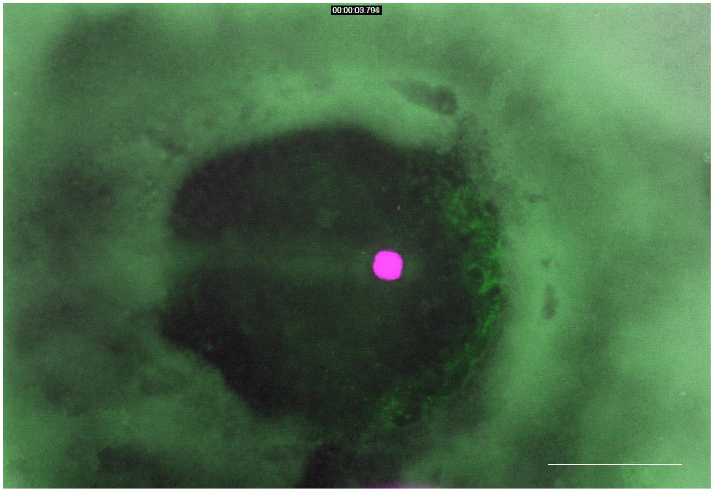
Development of mCherry-labeled Japanese quail node in homotopically grafted SN-EGFP-labeled chicken embryo in Supplementary Figure 3. (Frame 1)

**Movies G1 to G4.**
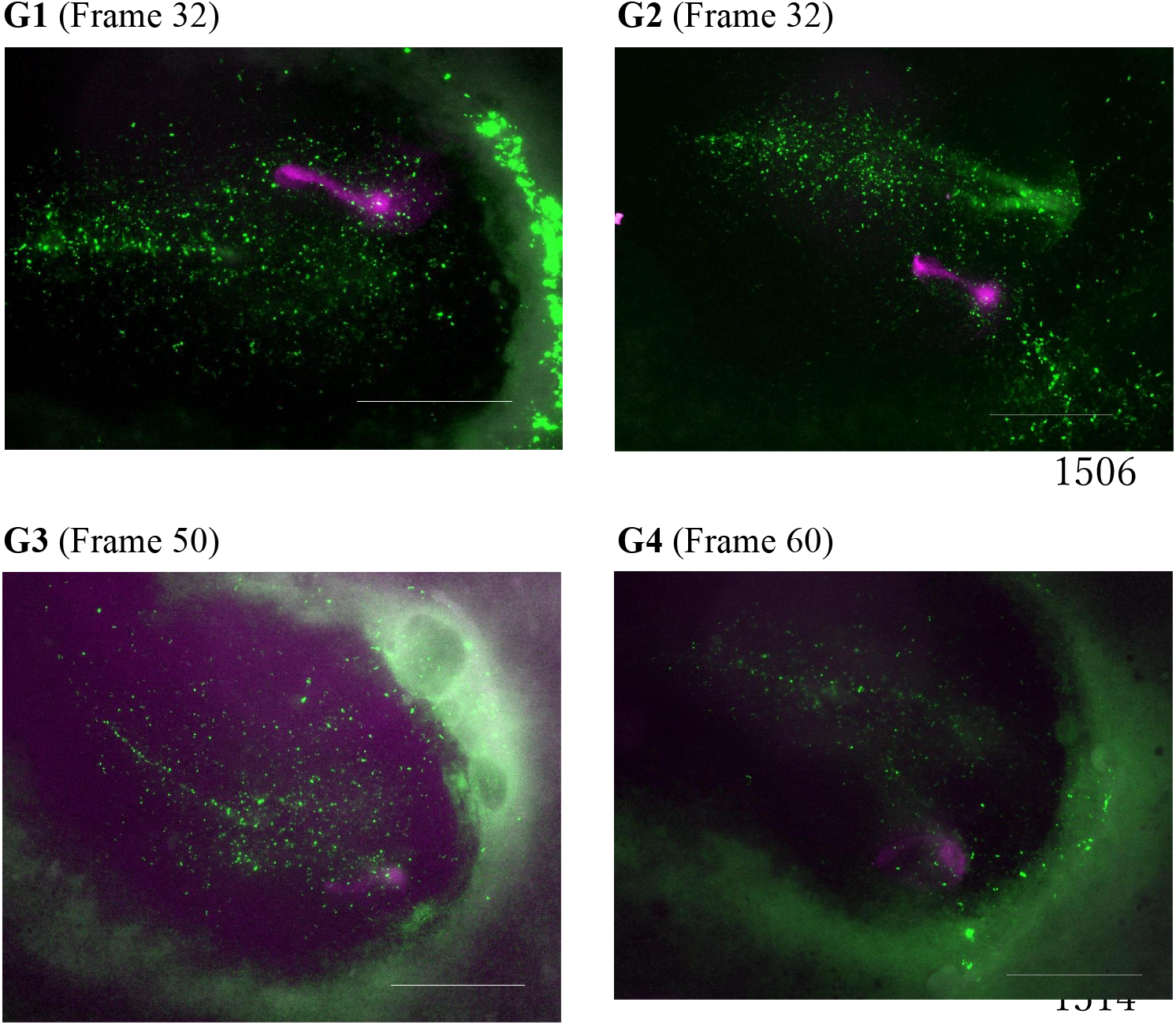
Original data of Fig. 7(A) to (D).

**Supplementary Table.** Correlations among the Figures, Supplementary Figures, and Movies

| Figures | Supplementary Figures | Movies |
| --- | --- | --- |
| Figure 1. The performance of Supernova-based epiblast cell labeling with EGFP. |  | Movie A. Original data of Figure 1(B). |
| Figure 2. Cell tracking-based analyses of collective cell migrations that form the head tissues. |  | Movie B1. Tracking SN-EGFP-labeled cells in Fig. 2(A)(a). |
|  |  | Movie B2. Tracking SN-EGFP-labeled cells in Fig. 2(A)(b). |
|  |  | Movie C1. Cell migration vectors in Fig. 2(B)(a). |
|  |  | Movie C2. Cell migration vectors in Fig. 2(B)(b). |
| Figure 3. Tracking the origin of head tissue portions at different axial levels back to st. 5. | Supplementary Figure 1. Additional cell tracking data between st. 4 and st. 6. | Movie D1. Backtracking data for Fig. 3(C). |
|  |  | Movie D2. The second example of backtracking data. |
| Figure 4. Comparison of the distribution of head tissue precursors at st. 4/5. |  | Movie E. Comparison of N2 and SN-EGFP for Figure 4(D). |
| Figure 5. Development of st. 4 nodes solely into the anterior mesendoderm and posterior notochord. | Supplementary Figure 2. The procedure to graft a Japanese quail node/AME in a st. 4 chicken host embryo. |  |
|  | Supplementary Figure 3. Development of mCherry-labeled Japanese quail node in homotopically grafted SN-EGFP-labeled chicken embryo. | Movie F. Development of mCherry-labeled Japanese quail node in homotopically grafted SN-EGFP-labeled chicken embryo in Supplementary Figure 3. |
| Figure 6. AP level-dependent development of the st. 4 quail node graft and host-derived secondary brain portions. | Supplementary Figure 4. The outcome of st. 5 node grafts at various anteroposterior levels of st. 4 embryos. |  |
| Figure 7. Live imaging of the events after grafting mCherry-transgenic quail node/AME in SN-EGFP-labeled st. 4 chicken embryos. | Supplementary Figure 5. Confirmation of head precursor convergence on gAMEs by cell track analysis. | Movie G1. Original data of Fig. 7(A). |
|  |  | Movie G2. Original data of Fig. 7(B). |
|  |  | Movie G3. Original data of Fig. 7(C). |
|  |  | Movie G4. Original data of Fig. 7(D). |
|  | Supplementary Figure 6. Detection of S-phase nuclei by EdU labeling of node-grafted epiblasts. |  |
| Figure 8. The gAME position-dependent development of the secondary brain portions. |  |  |
| Figure 9. The model of anterior epiblast regionalities with the potential to develop into FB, MB, and HB portions or abutting head ectoderm of the equivalent AP levels. |  |  |

## Notes

### Competing Interest Statement

The authors have declared no competing interest.

